# Description of Cranial Elements and Ontogenetic Change within *Tropidolaemus wagleri* (Serpentes: Crotalinae)

**DOI:** 10.1101/438093

**Authors:** Nicolette Hill

## Abstract

*Tropidolaemus wagleri* is a species of Asian pitviper with a geographic range including Thailand, Vietnam, Malaysia, Singapore, Bruniei, parts of Indonesia, and the Philippines. *Tropidolaemus* is a member of the Crotalinae subfamily, within Viperidae. The genus *Tropidolaemus* includes five species, and was once included within the genus *Trimeresurus*. While some osteologic characteristics have been noted a comprehensive description of cranial elements has not been produced for *T. wagleri*. An in-depth description of the cranial skeleton of *Tropidolaemus wagleri* lays the foundation for future projects to compare and contrast other taxa within Crotalinae and Viperidae. The chosen reference specimen was compared to the presumed younger specimens to note any variation in ontogeny. The study here provides a comprehensive description of isolated cranial elements as well as a description of ontogenetic change within the specimens observed. This study contributes to the knowledge of osteological characters in *T. wagleri* and provides a foundation for a long term project to identify isolated elements in the fossil record.

---

In order to best understand and evaluate the relationship between snake genera one must first understand the morphological variation observed within a single species. The in-depth description here of select cranial bones of *Tropidoleamus wagleri* (Wagler’s Palm Viper, Wagler’s Pitviper) is intended to act as a foundation to be used to compare and contrast other related species along with possibly identifying fossils recorded in Southeast Asia deposits. After this ‘base’ was created, ontogenetic change within *Tropidoleamus wagleri* was noted using morphological characteristics of the skull.

*Tropidolaemus* is a member of the Crotalinae subfamily within the family Viperidae (Reptilia: Squamata). Members of the Crotalinae possess a heat sensitive pit between the eye and the naris located on each side of the head (“pitvipers”). *Tropidolaemus wagleri* specimens often show laterally compressed bodies, slender in males while thick and stout in females. The laterally compressed body is a specialization for their arboreal lifestyle. *Tropidolaemus wagleri* has a geographic range including Southern Thailand, West Malaysia, Sumatra, Nias, Mentawei Archipelago, and Bangka Island (but not Belitung) [1]. They are typically found in wet, low elevation forests on lower branches of trees, and are viviparous [2]. The head shows a distinct triangular shape. Females display a black and yellow “speckled” coloration, their body is a glossy black with 25-30 irregular bright yellow vertical crossbars [1]. Males display a bright or deep green color with dorsolateral dots ranging from red at the anterior to white at the posterior [3]. *Tropidolaemus* exhibits dramatic ontogenetic color change and dramatic sexual dimorphism, further complicating the arrangement of species and possible subspecies [3].

*Tropidolaemus wagleri* possesses a complex taxonomic background. The genus is attributed to Wagler in 1830 according to McDiarmid et al. [4], Gumprecht et al. [5], and Vogel et al. [1]. There has been a long running discussion as to whom the name *T. wagleri* should be attributed to; H. Boie in 1826 (not seen), F. Boie in 1827 (not seen), or Schlegel in 1826 or 1827. Schlegel [6, 7] cited the name to H. Boie, whose manuscript contained descriptions, but was unpublished and therefore not considered valid. Schlegel [6, 7] did not include a description for the name, thus he cannot be correctly credited with the species name [1, 4, 5]. In most recent literature the species has been attributed to Boie 1827 (not seen), where he listed it under the genus *Cophias* [1, 4, 5]. While Boie did not provide a description, he did reference a previous description and definition in Seba [8], allowing for a description via indication [1, 4]. In 1830 Wagler reassigned the species to the genus *Tropidolaemus*, and then Brattstrom [9] regarded *Tropidolaemus* as a subgenus of *Trimeresurus*. After many synonyms and reassignments, Burger [10] resurrected the genus based on morphological characteristics and it has since been considered valid and supported by molecular studies [11, 12, 13.]. Hoser [14] suggested that a separate subfamily, Tropidolaemusiinae and a tribe Tropidolaemusini be created. Because these levels do not seem to be well founded, his taxonomy will not be followed in this paper. *Tropidolaemus* was initially accepted as monotypic, with the only species being *T. wagleri* until 1998 when *Trimeresurus huttoni* was shown to be a member of the genus [David and Vogel in Vogel et al. 1]. Vogel et al. [1] selected a neotype of *Tropidolaemus wagleri* and identified three additional species within the genus, for a current total of five species: *T. wagleri*, *T. huttoni*, *T. laticinctus*, *T. subannulatus*, and *T. philippensis*.

*Tropidolaemus spp.* are characterized by: “absence of a nasal pore, upper surfaces of the snout and head covered with distinctly keeled small scales, strongly keeled gular scales, second supralabial not bordering the anterior margin of the loreal pit and topped by a prefoveal, and a green coloration in juveniles which may or may not change with growth” [1 p2]. Following Vogel et al. [1 p2] *Tropidolaemus wagleri* is characterized by:

- Internasal scales always touching
- Background body color with clear ontogenetic variation: black (never green) in adult females, whereas males and juveniles retain a vividly green background color
- Pattern with clear ontogenetic variation: yellow crossbands around the body in adult females, white spots in adult and juvenile males, and white crossbars in juvenile females
- Postocular stripe with ontogenetic variation: black stripe in adult females whereas males and juveniles of both sexes exhibit a white and red stripe
- Belly pattern: banded in adult females while uniform in males and juveniles
- Number and keeling of dorsal scale rows at midbody: feebly keeled with 21–23 in males and distinctly keeled with 23–27 in females
- Ventral plates: 143–152 in males and 134–147 in females
- Subcaudal plates: 50–55 in males and 45–54 in females

Descriptions of the cranial skeletal elements of select taxa within Viperidae have previously been constructed by many authors. Dullenmeijer [15] described the head of the common viper (*Vipera berus*, Viperinae), with functional morphology aspects. Brattstrom [9] looked at pitviper osteology to better understand the evolution of the group. He described multiple species with hand-drawn illustrations of the various bones, and described the importance of certain osteologic characters of *Tropidolaemus wagleri*. Burger [10] looked at multiple pitviper genera and their characteristics. Lombard [16] looked at the shape of the ectopterygoid of Colubroidea using morphometrics. Guo et al. [17] compared skull morphology of Crotalinae species. Guo and Zhao [18] aimed to identify diagnostic characters for some of the genera within *Trimeresurus* (sensu lato). A number of characters were found to be important in distinguishing species and genera such as the size of the ectopterygoid anterior lateral process, the shape of the palatine, and the shape of the frontal. Cundall and Irish [19] described characteristics of several different families of snakes. Guo et al. [20] examined the systematic value of using skull morphology to distinguish between members in the *Trimeresurus* radiation. Different characters exhibited various success within the different pitvipers studied, leading them to conclude that “skull morphology can contribute to an overall understanding of pitviper taxonomy, but that it would be unwise to rely on skull characteristics alone” [20 p378]. Guo et al. [21] observed the evolution of 12 osteological characters in 31 species of Asian pitvipers in order to form a phylogenetic perspective.

Due to *T. wagleri* being one of the first Asian pitvipers described, and its substantial sexual dimorphism and geographic and ontogenetic variation, many specimens were initially identified as other species [1, 11]. Within tropical Asia pitvipers of the genus *Tropidolaemus* are the most widespread and commonly encountered venomous snakes in many islands of the Malay Archipelago [1]. The wide geographic distribution and prevalence of human interaction demonstrate the importance of understanding *T. wagleri*. The descriptions in this study have the potential to contribute to many future projects spanning a variation of emphases.

## Methods and results

Multiple specimens of the same age were located to begin the description of *Tropidoleamus wagleri*. Specimens were compared to ensure the characteristics are consistent within each specimen of *T. wagleri*. Twelve cranial elements were studied: quadrate, ectopterygoid, pterygoid, palatine, maxilla, parietal, prefrontal, premaxilla, dentary, angular, splenial, and compound bone. Elements chosen were expected to show the most ontogenetic variation, and are often distinct between snake taxa. Among previous works on pitviper cranial osteology, Guo and Zhao [18] presented diagnostic characters for nine Asian pitvipers (not including *Tropidolaemus*); their characters include the shape of the palatine, shape of the frontal, presence or absence of a projection on the border of the pit cavity, and the size of the ectopterygoid anterior lateral process. These and other characters were recorded for *T. wagleri* specimens in the study presented here. All distinct characters are described in detail allowing for future in-depth comparisons to other species. After the descriptions were produced, the skeletal elements were then compared to those in other *T. wagleri* specimens of different ages. The distinct characters used to describe *T. wagleri* were used as a reference for any changes through ontogeny.

Twelve specimens were used for this study, 11 from the East Tennessee State University Vertebrate Paleontology Collections (ETVP) and one from Appalachian State University (ASU). In addition, an articulated specimen from Appalachian State University (APPSU 25497) was used as a guide to an articulated skull but not used for descriptions (Figs. 1-3). Of the 12 specimens, only one (ASU 19452) had a recorded snout-vent length (SVL). Burger [10] reported a total length of 700mm while Kuch et al. [3] gave a range of 350–1000mm for total length. One study reported a maximum SVL of 770mm for a female and 435mm for a male [1]. This study functions under the assumption that greater size is equivalent to greater age. In order to determine the relative age of the remaining 11 specimens the SVL had to be calculated. Length of the quadrate was used as a proxy to estimate a snout-vent length. Using quadrate length to estimate SVL, the 12 specimens were then arranged by size and categorized into an age rank (AR) series based on estimated SVL; where 1 is the youngest (neonate) and 12 is the oldest. Quadrates were measured along the ventral side, creating a greatest anteroposterior length.

**Fig. 1.**
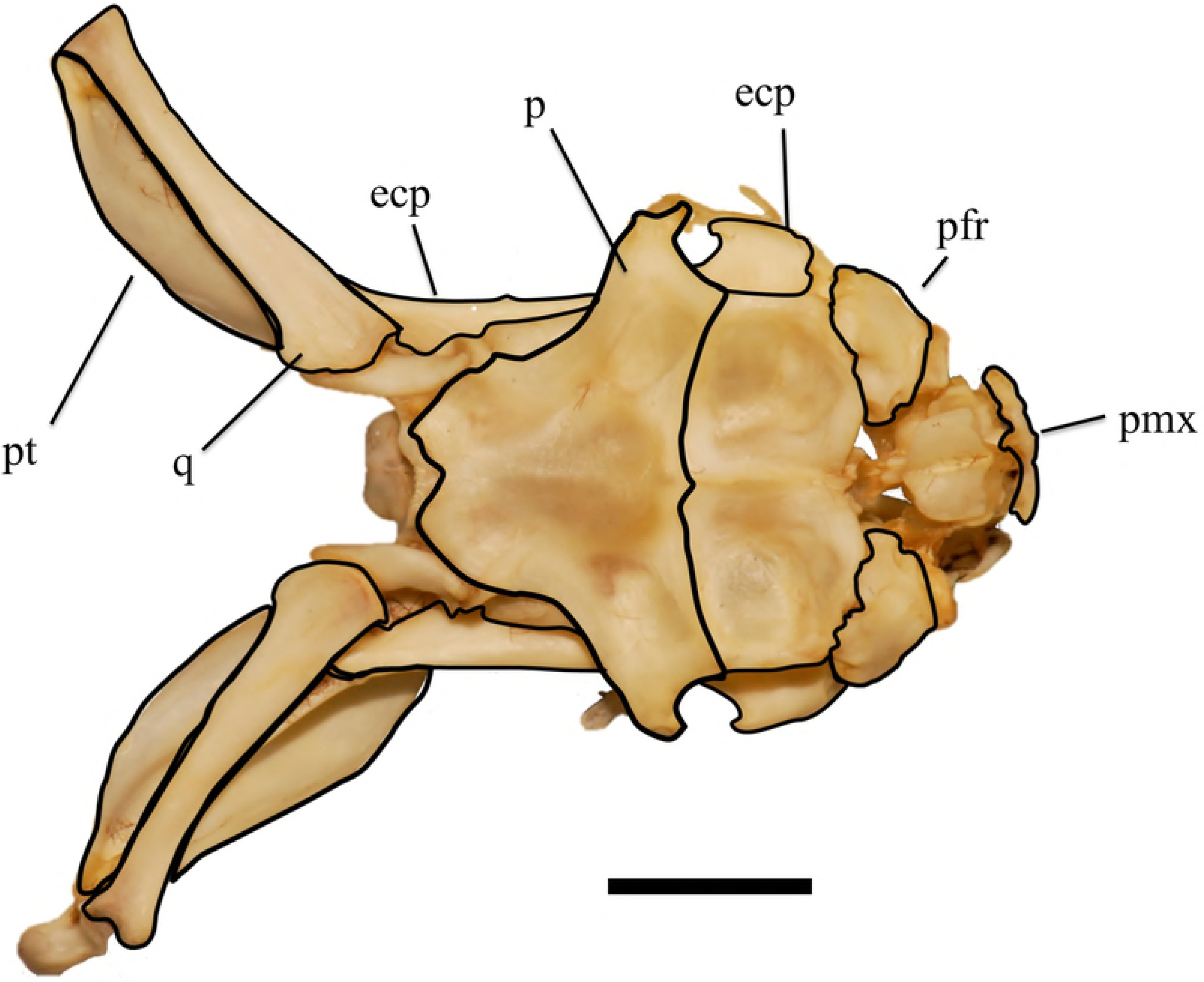
Skull of *Tropidolaemus wagleri* (APPSU 25497) in dorsal view. Anterior is to the right. Scale bar = 10 mm. Abbreviations: ecp = ectopterygoid; p = parietal; pfr = prefrontal; pmx = premaxilla; pt = pterygoid; q = quadrate.

**Fig. 2.**
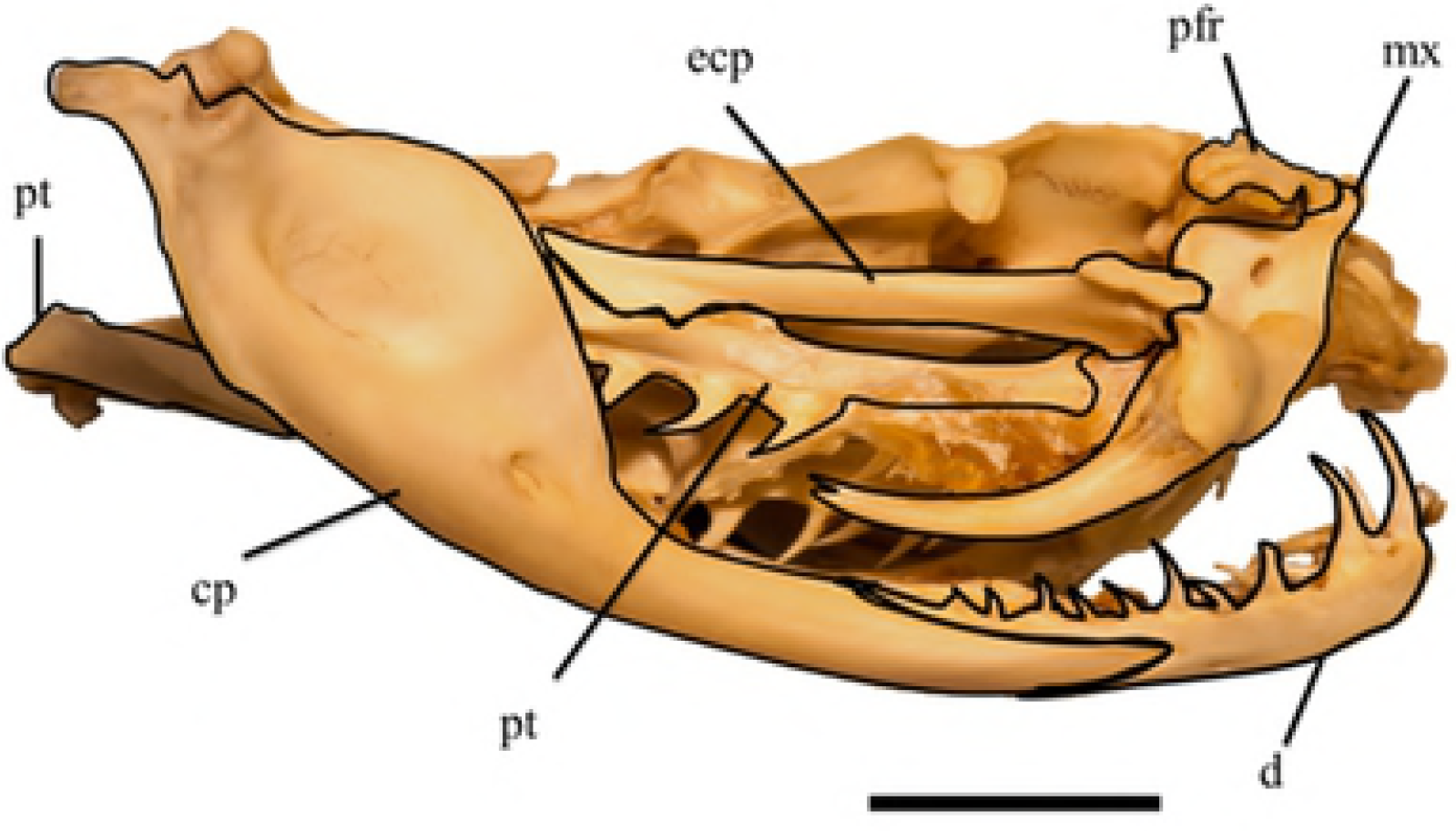
Skull of *Tropidolaemus wagleri* (APPSU 25797) in right lateral view. Anterior is to the right. Scale bar = 10 mm. Abbreviations: ecp = ectopterygoid; cp = compound bone; d = dentary; mx = maxilla; pfr = prefrontal; pt = pterygoid.

**Fig. 3.**
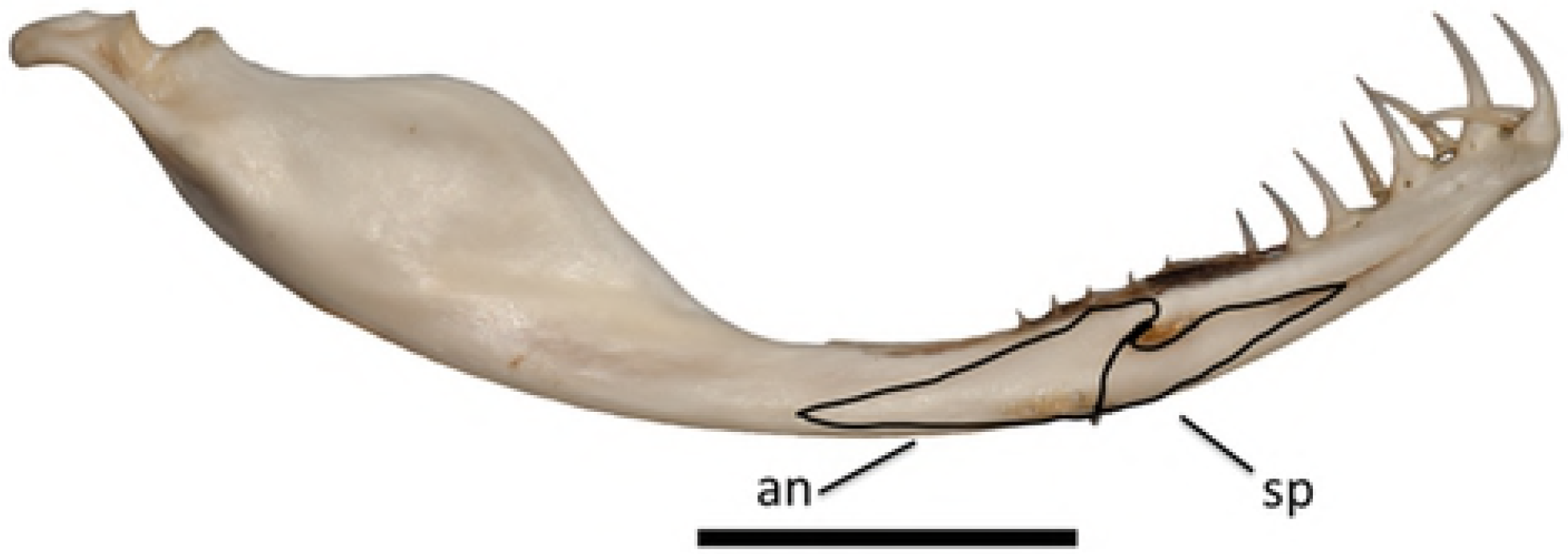
Lower left jaw of *Tropidolaemus wagleri* (APPSU 25797) in medial view. Anterior is to the right. Scale bar = 10 mm. Abbreviations: an = angular; sp = splenial.

The quadrate of specimen ASU 19452 was measured, and the quadrate-to-SVL ratio was then used to calculate the SVL for the rest of the specimens (Table 1). A total of 12 elements were studied and described, only using the right of paired elements, following the terminology used in Cundall and Irish [19] and Evans [24]. Abbreviations from Cundall and Irish [19] were followed. When not available in Cundall and Irish (19) abbreviations from Evans [24] were followed. When neither were available abbreviations were created. The chosen reference specimen (ETVP 3295) was then compared to other age ranks in order to record any ontogenetic changes. Reference specimen was chosen due to it being the largest specimen and therefore, the presumed most mature.

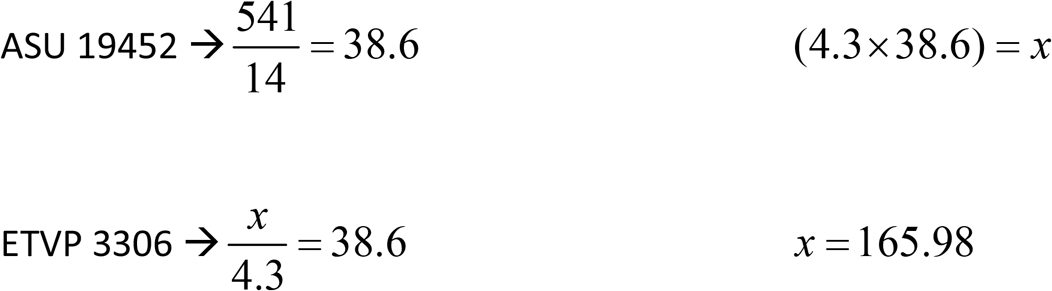

**Table 1.** Calculated SVL (mm) and Age Ranks (AR) using quadrate length (mm). Reference specimen is indicated by asterisk*

| Age Rank | Specimen ID | Quadrate length (mm) | Calculated SVL |
| --- | --- | --- | --- |
| 1 | ETVP 3306 | 4.3 | 166 |
| 2 | ETVP 3290 | 13.0 | 502 |
| 2 | ETVP 3293 | 13.0 | 502 |
| 3 | ETVP 3298 | 13.5 | 521 |
| 4 | ASU 19452 | 14.0 | 541 |
| 5 | ETVP 3296 | 14.3 | 552 |
| 6 | ETVP 3292 | 15.0 | 579 |
| 7 | ETVP 3300 | 15.3 | 591 |
| 8 | ETVP 3301 | 18.0 | 695 |
| 9 | ETVP 3297 | 18.5 | 714 |
| 10 | ETVP 3294 | 19.0 | 734 |
| 11 | ETVP 3299 | 19.5 | 753 |
| 12 | ETVP 3295* | 20.0 | 772 |

## Descriptions

### Quadrate

The anterior end of the quadrate overlaps the posterior of the supratemporal. Posterior end forms a mandibular condyle that interlocks with the articular of the compound bone. The posteromedial end adjoins the posterior of the pterygoid.

#### Dorsal

Anterior end is slightly curved with a flat edge. The conch is also located at the anterior end, is concave, and spans half the length of the bone. The tips of the mandibular condyle gently flare medially. Distally it has a large curve forming a process that curves into the mandibular condyle (Fig 4).

#### Ventral

Anterior end is the widest part of the bone. It is moderately curved with a flat edge. No conch is present. Less than half way down the shaft, the articulatory process of the quadrate is present. This articulatory process is distal in ventral view on the right quadrate, and has a flat surface. It articulates with the columellar cartilage. Posterior end shows a larger part of the mandibular condyle. Medial side curves outward significantly. Distal end appears to extend slightly past the articulation surface. The articulation surface appears grainy compared to the smooth bone of the shaft. There appears to be a small fossa close to the distal end

#### Ontogeny

Anterior, spatulate end has a rough, textured edge in AR1; in AR2 the edge is smoother, but still more rough than in AR12. The anterior end is more flared and widens quickly in AR1, while the widening occurs more gradually in AR2, much like in AR12. Articular process on the ventral side protrudes more and is slightly more elongate in AR2. Lateral process of the mandibular condyle is more rounded and protrudes greater in AR1. The notch is also not completely formed in AR1, medial side of the lateral process curves back anteriorly to meet the shaft, there is no process medially.

##### Summary of Ontogeny

- Anterior margin smoothes in texture with age.
- Anterior half “fills out” proportionally with age; widening appearing as a dramatic flare in AR1 and ending as a gradual widening in AR12.
- Lateral process of mandibular condyle widens and recedes closer to the bone with age; appearing more knob-like rather than a drastic projection.
- Articular surface of the mandibular notch exhibits less dramatic projections with age.

**Fig. 4.**
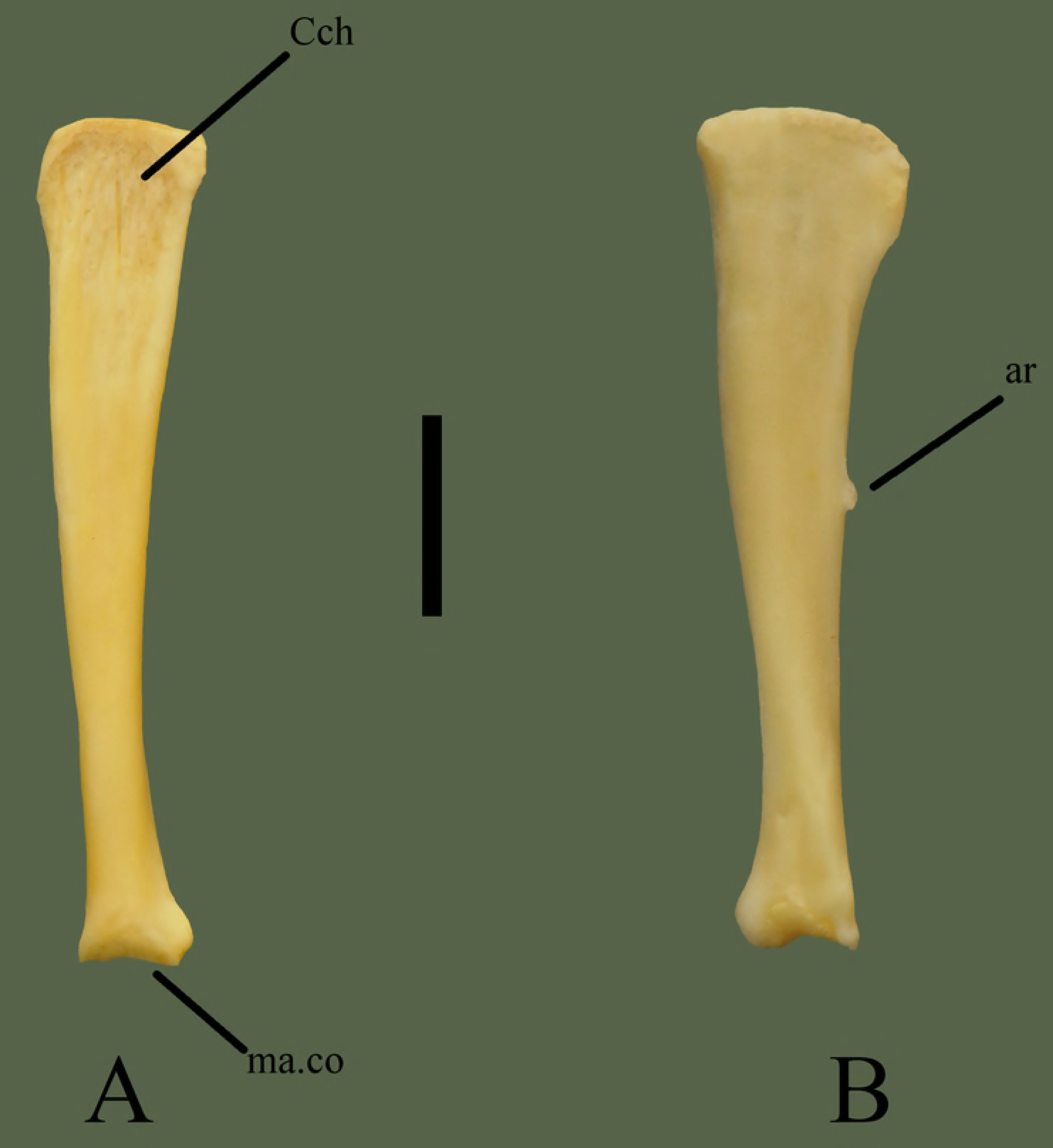
Right quadrate (ETVP 3295) in dorsal (A) and ventral (B) view. Anterior is to the top. Scale bar = 5 mm. Abbreviations: ar = articulatory process of quadrate; Cch = conch; ma.co = mandibular condyle.

### Ectopterygoid

Anterior lateral and ventral processes of the ectopterygoid cup the lateral facet located on the posterior side of the transverse crest, dorsal to the dentigerous region of the maxilla. The anterior medial process of the ectopterygoid fits into the posteromedial facet of the transverse crest of the maxilla. Posterior end of the ectopterygoid overlaps the pterygoid in the middle of its lateral curvature, with the posterior ventral process of the ectopterygoid contacting the lateral process of the pterygoid. The thin, angled portion of the posterior of the ectopterygoid fits into the groove on the dorsal side of the pterygoid.

#### Dorsal

Anterior end is bifurcated, producing two processes, one medial and one lateral. The lateral process is anteroposteriorly broad and is not as pronounced as the medial process. Lateral process is rectangular in shape, with a point at the posterior end that curves medially back toward the shaft of the bone. Medial process is narrow and protrudes anteromedially from the bone. Between the two anterior processes is a notch where the maxilla articulates. The shaft is predominantly straight and flat; most modification is present at the ends. Posterior end angles medially to allow for articulation with the pterygoid. Directly anterior to the tip, on the medial side, there is the small pterygoid process that permits articulation and kinesis with the ectopterygoid (Fig 5).

#### Ventral

Lateral process curves into itself on the ventral side. Lateral to the notch is a rounded process, this process articulates with the short ridge on the dorsal side of the pterygoid. The medial process does not show s difference in ventral view. Approximately 13 mm posterior from the most posterior point of the lateral process a fossa is present. The fossa exhibits a crest on the inferior side that curves dorsally into a short ridge that terminates ~3 mm before the posterior end. Posterior end curves out laterally to terminate at a rounded, laterally angled process. Medial side of the terminating process curves in medially ~5 mm to a point. The point is the posterior edge of a flat, rectangular shaped process. Rectangular process is ~2 mm long and the anterior point curves laterally to meet the shaft of the ectopterygoid. A short ridge is visible on the medial side, beginning at the center of the rectangular process and terminating ~4 mm anterior of the process.

#### Ontogeny

Medial process is distinctly thin in AR1, comes to point. In AR2 the shape is formed, thicker, but still thinner than in AR12. Small medial process on AR2, posterior to main process, not present in others. In AR1 posterior end comes to sharp point. In AR2 the shape of posterior is formed, medial side of posterior does not exhibit the flat protrusion like in AR12. The flat protrusion is still not form in AR10 and AR11. Hooked shape of lateral process slowly forms from small to large. In AR1 the anterior of the lateral process consists of two processes, in AR12 and AR2 the lateral process appears as one rounded process. Ventral process present in AR12 and AR2, AR1 has a short ridge from medial process that terminates where ventral process will form. Posterior end comes to a point in AR1, concave with higher sides. The ventral process of the posterior end not yet formed in AR1. Medial margin remains straight to medial process in AR1. In AR2 the medial side curves to pointed process then curves to medial process. In AR12 the medial side gently curves to the medial process with no extra processes.

##### Summary of Ontogeny

- Medial process forms distinct shape with age; beginning as a thin, pointed process and forming into thick, square-shaped process.
- Posterior tip thickens with age and flat protrusion forms with age.
- Lateral process begins with an additional process on anterodorsal side and morphs into one rounded process with age.
- Anterior ventral process forms between AR1 and AR2.
- Posterior end forms distinct shape with age.

**Fig. 5.**
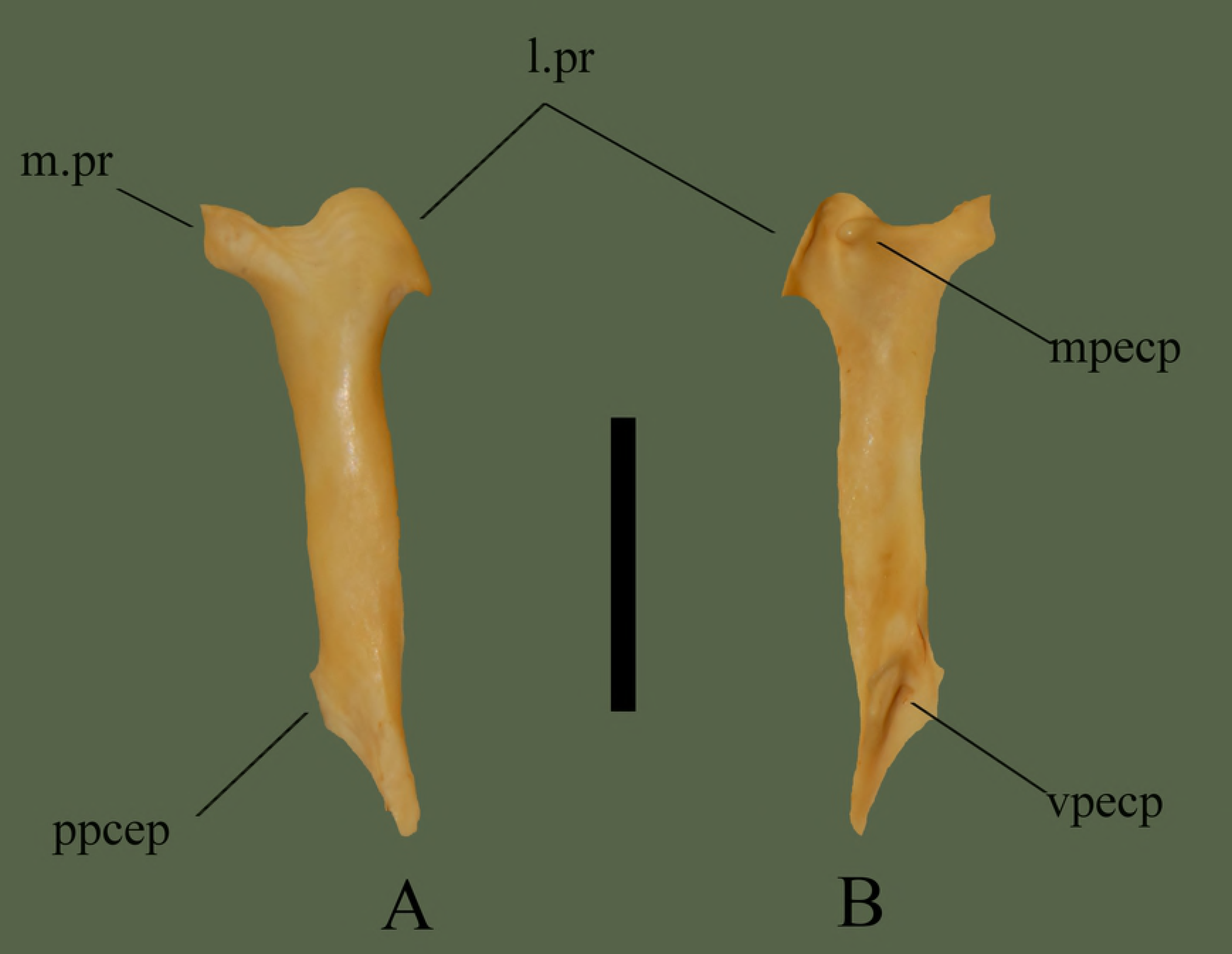
Right ectopterygoid (ETVP 3295) in dorsal (A) and ventral (B) view. Anterior is to the top. Scale bar = 10 mm. Abbreviations: l.pr = lateral process; m.pr = medial process; mpecp = maxillary process of ectopterygoid; ppecp = pterygoid process of ectopterygoid; vpecp = ventral process of ectopterygoid.

### Pterygoid

The curved anterior tip and process of the pterygoid interlocks between the dorsal and ventral processes of the palatine, the posterolateral end sits against the medial side of the quadrate and borders the articular of the compound bone. Groove on the dorsal side of the pterygoid cups the posterior ventral process of the ectopterygoid.

#### Dorsal

The pterygoid has a dentigerous region that makes up a little more than half the length of the bone. Dentigerous region is long and narrow, making up the anterior half of the bone. Anterior tip exhibits undulations with a prominent anterior process marginally oriented laterally. About half way down the bone a process begins to form by the flange widening the dentigerous region medially. The process is laterally oriented and comes to a point. Located posteromedially to the lateral process, on the body of the bone is a short ridge with an associated groove on its lateral side. The groove receives the posterior most of the ectopterygoid and the ridge fits into a complimentary groove. Posterior section of the bone consists of a broad flange that curves out medially then terminates in a rounded end angled laterally. Medial edge of the anterior end has a shallow groove that runs the length of the narrow dentigerous region. Groove widens at the posterior end and terminates before the widening of the dentigerous region. Posterior tip is rounded, with a wide groove that runs along the lateral side ~10 mm anteriorly. Inside the posterior groove, the terminating end appears to be folding in on itself. The area also exhibits two fossa, and has an overall folded texture (Fig 6).

#### Ventral

Anterior end curves medially to terminate at a narrow, rounded process. The pterygoid teeth are visible in ventral view, demonstrating their recurved teeth. Approximately 13 mm posterior from the anterior end the bone begins to widen laterally. The narrow dentigerous region remains distinct and angles medially terminating on the flange, ~9 mm before the posterior end. Widening comes to a point at ~3 mm; this pointed process is oriented laterally. Posterior to the pointed process the flange begins to curve out laterally, terminating at the medially angled posterior end. The end is rounded, and slightly oblong shaped. Medial edge of the posterior end curves out anteromedially for ~9 mm, and then curves in anterolaterally to meet the narrow dentigerous region. A wide, long groove is present, beginning at the point of widening and terminating at the point of inflection at the posterior end. On the lateral side of the flange a dorsally curved ridge runs ~11mm down the bone.

#### Ontogeny

In dorsal view a short ridge is visible down the center of the flange, beginning where the flange starts in AR1; this ridge might form the rounded dorsal process as seen on AR12 and AR2, though less developed. The edges of AR1 are rough rather than smooth as in AR2 and AR12. In ventral view, AR1 does not have a prominent projection where the dentigerous region meets the medially oriented flange. Lateral side of flange is flat in AR12, mostly flat in AR2 and has a groove in AR1. Teeth not present in AR1, but tooth attachment areas are visible. In ventral view of AR2 the lateral process is present but not as prominent as in AR12. Anterior dorsally oriented process extends further anteriorly with age.

##### Summary of Ontogeny

- Short ridge on dorsal side of flange in AR1 may form the rounded dorsal process seen in AR2 and AR12.
- Medial projection where dentigerous region meets flange becomes more prominent with age.
- Lateral side of flange flattens with age.
- Anterior dorsally oriented process extends anteriorly with age.

**Fig. 6.**
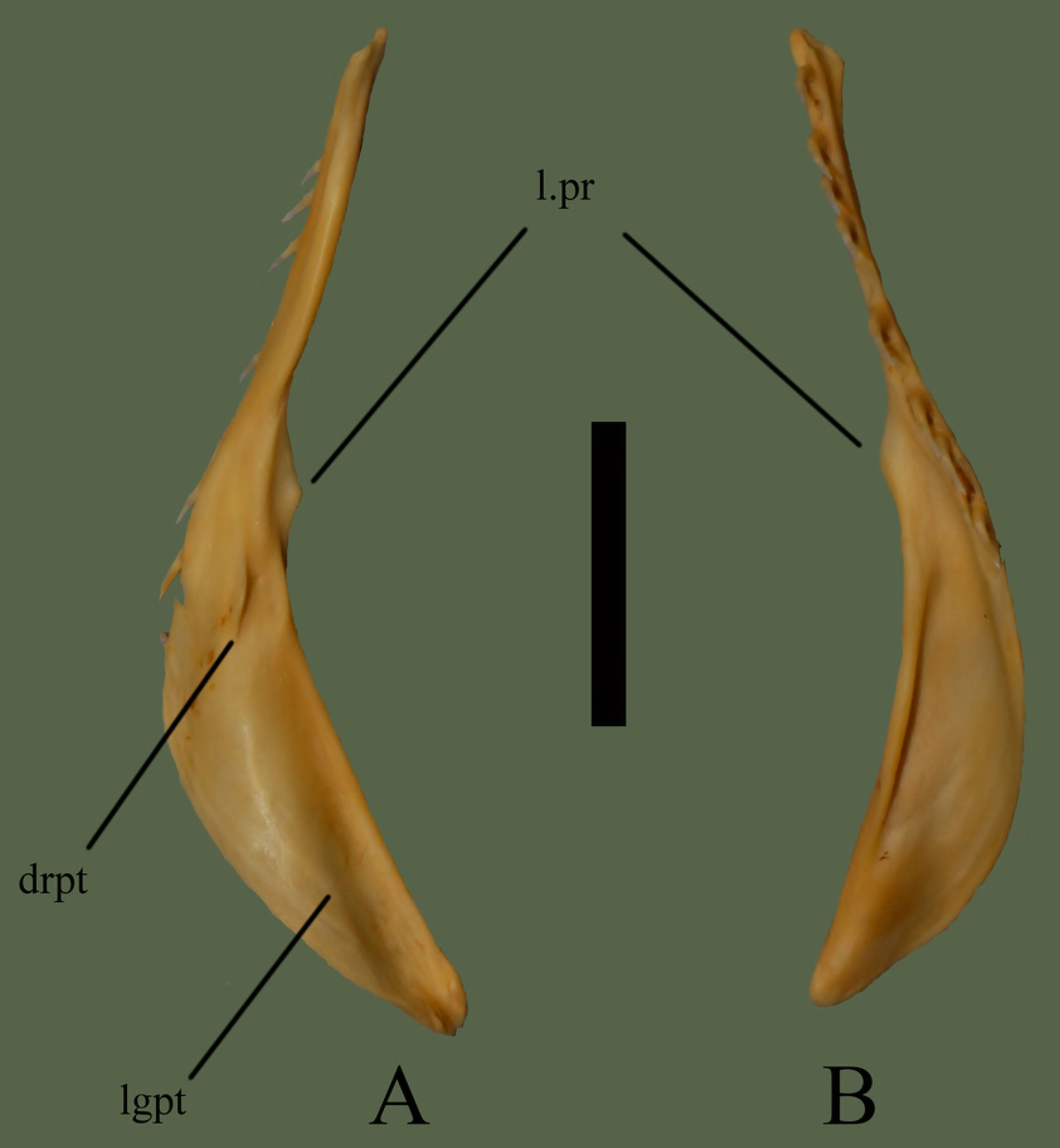
Right pterygoid (ETVP 3295) in dorsal (A) and ventral (B) view. Anterior is to the top. Scale bar = 10 mm. Abbreviations: l.pr = lateral process; drpt = dorsal ridge of pterygoid; lgpt = lateral groove of pterygoid.

### Palatine

Right palatine has three recurved teeth while the left has four. The posterior dorsal and ventral processes interlock with the curved anterior tip and anterior process of the pterygoid.

#### Medial

Anterior surface curves back posteriorly. A 1 mm section, directly dorsal to the first tooth, is less angled than the rest of the anterior end, appearing more flat. Medial side is slightly concave, with the posterior end near the articulation being more concave than the rest. Posterior end exhibits a forked dorsal and ventral process. These processes articulate with the anterior end of the pterygoid. The overall shape of the palatine is crescent-shaped [18]. Multiple fossa are present on the medial side (Fig 7).

#### Lateral

The lateral side is slightly convex with the dorsal side flaring out laterally. The palatine is wider ventrally, thinning as it terminates dorsally. Attachment area of each tooth extends past the base of the ventral border, forming dorsally oriented grooves between each tooth.

#### Ontogeny

AR2 has two teeth present with three, possibly four sockets. AR1 has no teeth and tooth attachment areas that may not yet be sockets, they are very narrow and very shallow. Dorsal side of AR1 is straight anteriorly and posteriorly curves ventrally. AR2 exhibits a thin anterior tip, curving up dorsally, forming a rounded dorsal edge and curving ventrally to terminate. The transition to a flat, rounded, dorsal edge is more extreme in AR2. The small posterior notch seen on AR12 appears to be forming in AR2; dorsal process is present, ventral process is oriented medially, not yet ventral. Posterior tip of AR1 shows a narrow groove separating two projections. One (dorsal) is rounded; the other (ventral) is thin, long, with a rounded tip. In medial view the two posterior processes of AR1 are more visible. One is thin and long while the other is short and thick. Posterior ventral process is present in AR2, it is thin and long, extending posteriorly more than in AR12.

##### Summary of Ontogeny

- Dorsal border fills out it’s shape with age.
- Posterior ventral process shortens and thickens with age
- Posterior notch forms with age

**Fig. 7.**
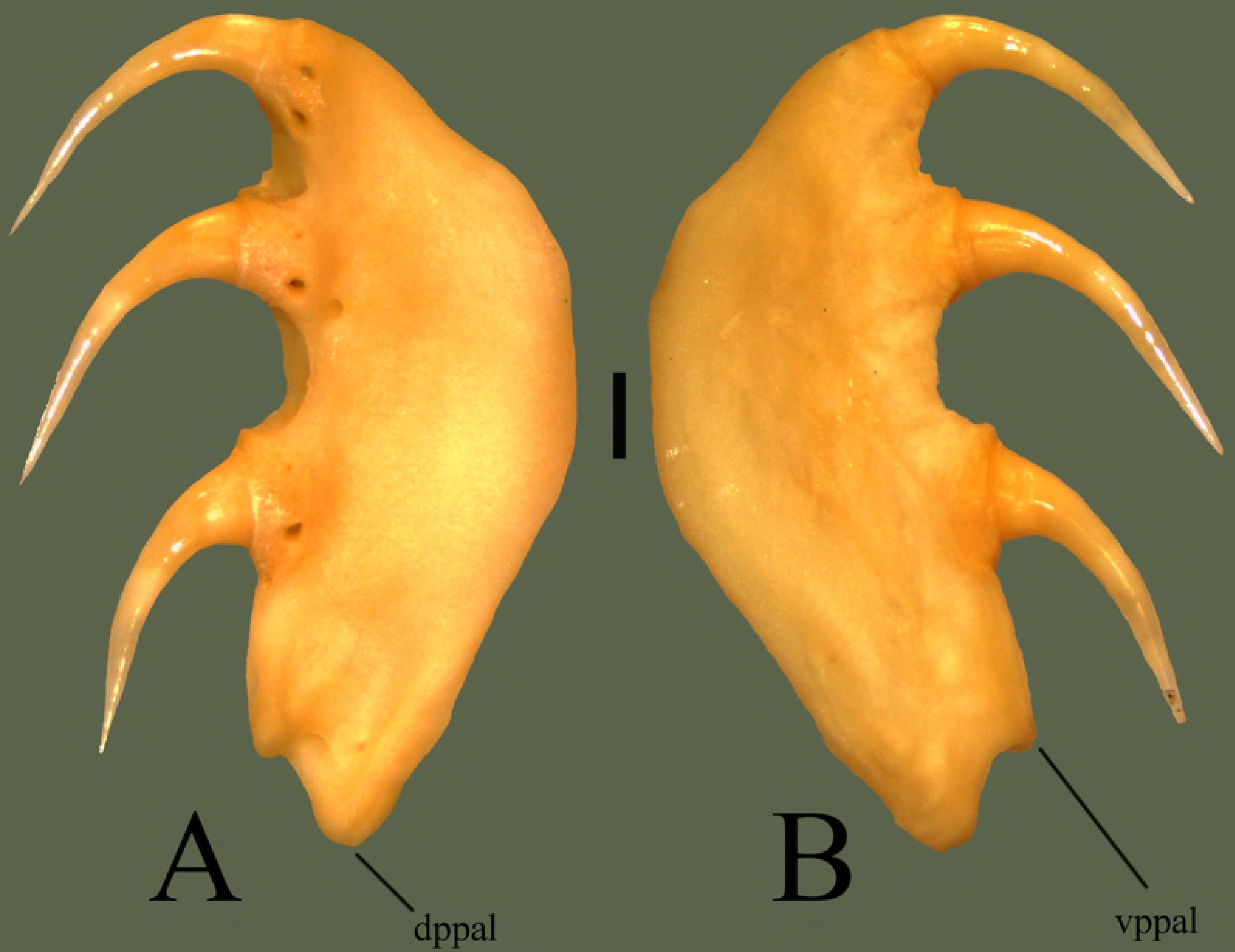
Right palatine (ETVP 3295) in medial (A) and lateral (B) view. Anterior is to the top. Scale bar = 1mm. Abbreviations: dppal = dorsal process of palatine; vppal = ventral process of palatine.

### Maxilla

The maxilla articulates posterodorsally to the prefrontal; the cupped articulation surface on the ascending process articulates with the maxillary facet on the anterior of the prefrontal. Posterior side of the transverse crest of the maxilla articulates with the anterior process of the ectopterygoid. Medial facet of the transverse crest receives the medial process of the ectopterygoid while the lateral facet of the transverse crest receives the lateral and ventral processes of the ectopterygoid. The maxilla does not articulate to the palatine, but is connected by cartilage.

#### Anterior

There is a dorsally oriented ascending process that meets the ventral surface of the prefrontal. Ventrally the ascending process curves laterally, forming a rounded process, then medially to meet the dentigerous base. The base is wide, mostly laterally. The ventral edge is textured and curves inward dorsally, bearing the fangs. The rounded process present on the anterior edge of the lateral opening was found to be taxonomically important in Brattstrom [9]. Guo and Zhao [18] defines two types of maxilla, type A which exhibits no projection, and type B which exhibits a projection on the anterior edge of the pit cavity. The maxilla observed for *Tropidolaemus wagleri* falls into type B (Fig 8).

#### Posterior

Dorsolateral to the fang is a transverse crest, when articulated it is located anterior to the edge of the ectopterygoid joint. Maxilla exhibits a large lateral opening, concave and extending anteromedially and dorsally from the transverse crest. Lateral opening holds the loreal pit. Dorsally oriented ascending process exhibits a cupped articular surface for attachment to the prefrontal. Posterior edge of the lateral opening is curved, forming two rounded processes. Lateral to the processes a long fossa is present. The fossa is longer than wide and contains an additional fossa.

#### Ontogeny

In anterior view the tip of the ascending process in thickest in AR12, and the body of the process is thickest in AR2 (Figs 9 and 10). Medial curve of the ascending process curves inward medially the most in AR12. Processes of the ascending process are thicker and exhibit greater curvature in AR12. In AR1 the ascending process is very thin with a gentle lateral curve. Lateral flange of the dentigerous region is present in AR1 but no major curvature has developed. A thickened area dorsal to the dentigerous region exhibits a lateral outward curvature, appearing to be the lateral process. In AR1 the posterior is visible in anterior view, curving outward laterally. Prefrontal facet of AR1 is more medially oriented and dorsoventrally longer than the others. In posterior view the prefrontal facet of AR12 exhibits a square shape. In AR2 the prefrontal facet is rounded while in AR1 it is laterally rounded and medially straight. Lateral process of ascending process is thinner and more prominent in AR12. It is rounded, thick, and short in AR2 and very short in AR1. Anterior covering of the dentigerous region is extremely short in AR1 while it is developed in both AR12 and AR2, with AR12 being most robust. Ascending process bears a fossa posteriorly; in AR2 it is thinner and dorsoventrally shorter than in AR12. In AR1 the fossa spans the majority of the length of the ascending process with the inner fossa accounting for half of the major fossa. Posterior ridge dorsal to dentigerous region is barely visible in AR1 and in AR2 it is thicker than in AR12.

##### Summary of Ontogeny

- Maxilla overall starts thin in AR1 and becomes more robust by AR2.
- Anterior covering of dentigerous region becomes more robust with age.
- Shape and curvature of ascending process develops with age.
- Posterior fossa minimizes, moves medially, and shallows from AR1 to AR2, then deepens and elongates dorsoventrally in AR12.

**Fig. 8.**
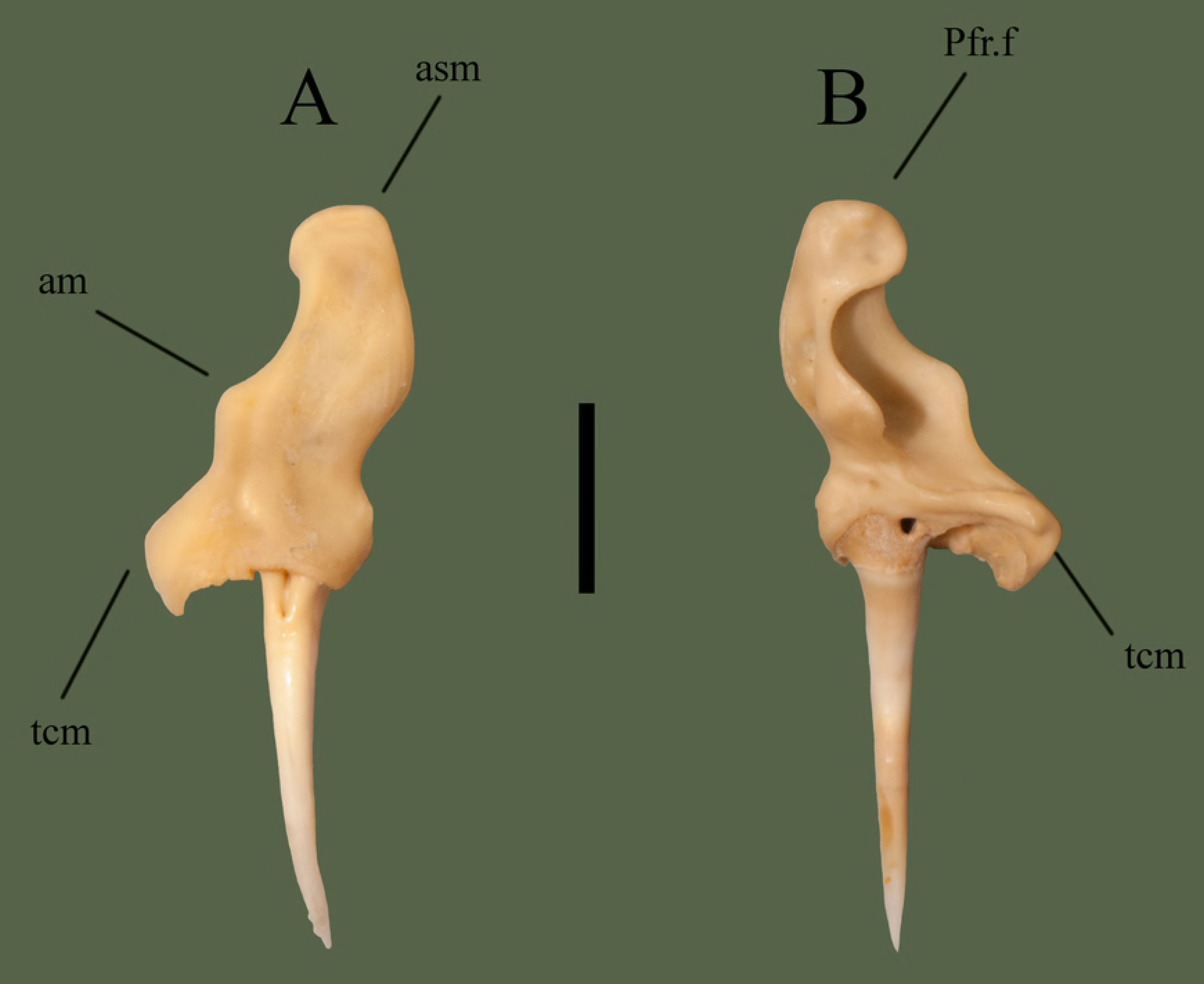
Right maxilla (ETVP 3295) in anterior (A) and posterior (B). Scale bar = 5 mm. Abbreviations: Pfr.f = prefrontal facet; am = anterior process of maxilla; asm = ascending process of maxilla; tcm = transverse crest of maxilla.

**Fig. 9.**
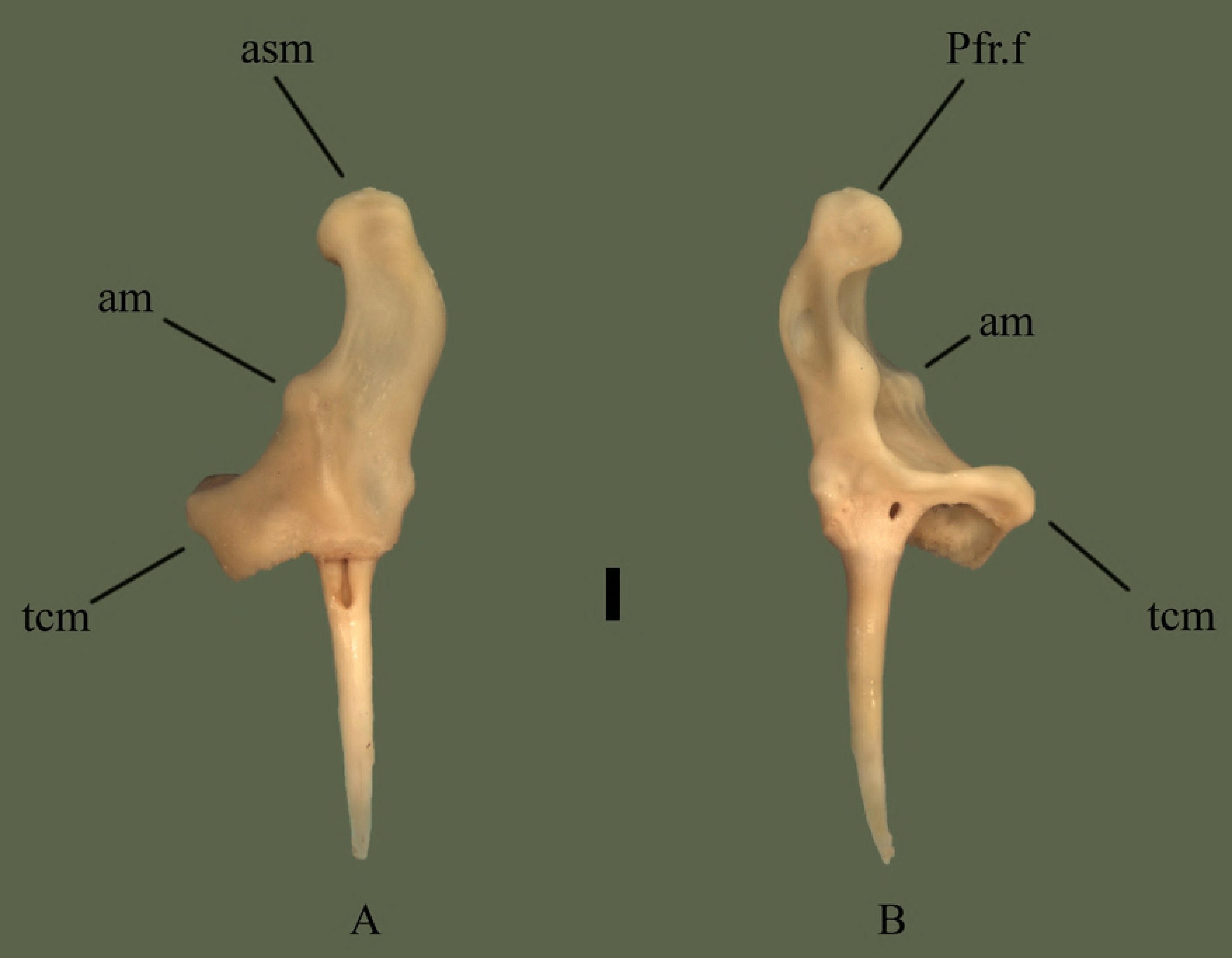
Right maxilla (ETVP 3306) in anterior (A) and posterior (B). Scale bar = 1mm. Abbreviations: Pfr.f = prefrontal facet; am = anterior process of maxilla; asm = ascending process of maxilla; tcm = transverse crest of maxilla.

**Fig. 10.**
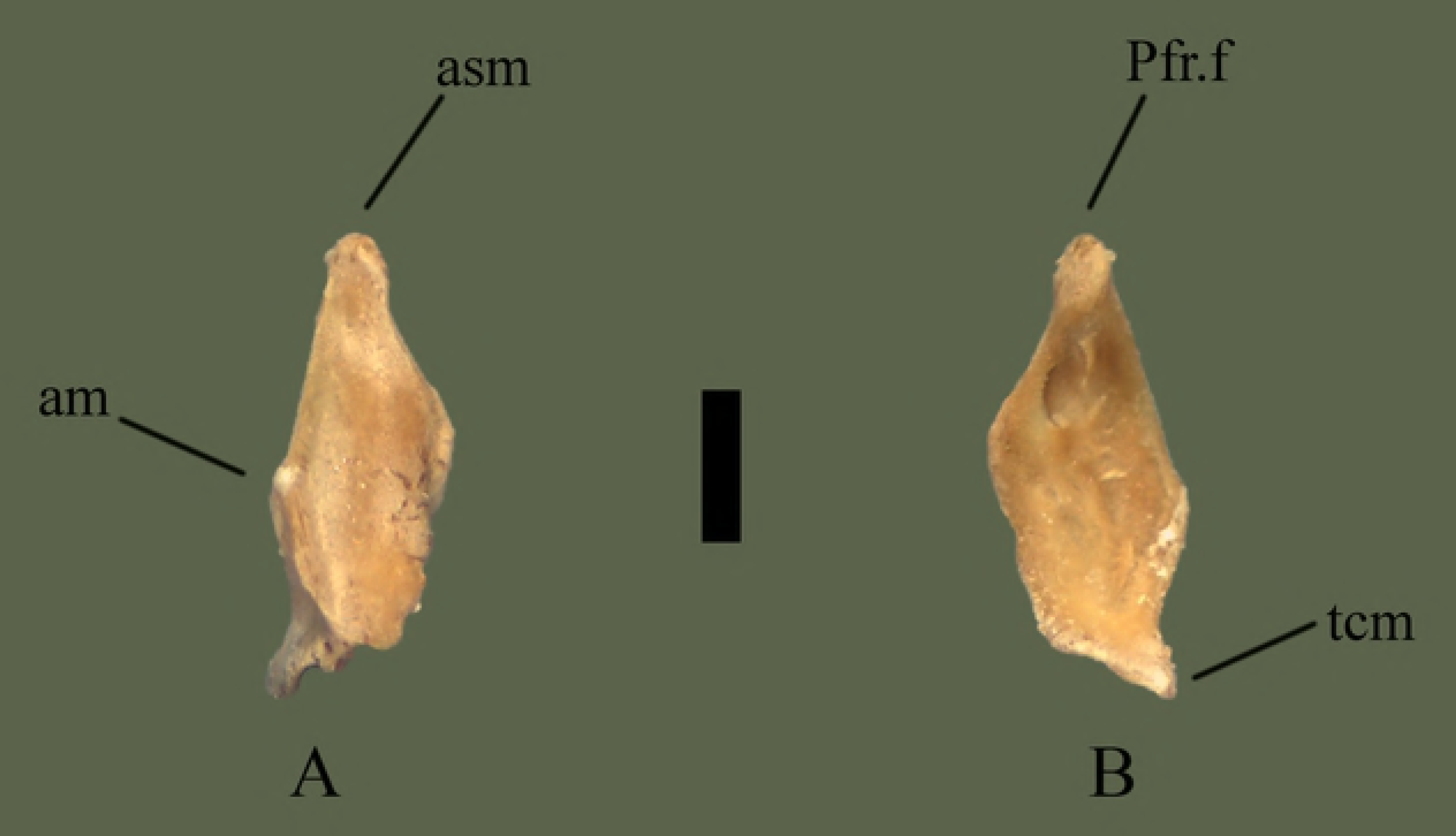
Right maxilla (ETVP 3306) in anterior (A) and posterior (B). Scale bar = 1mm. Abbreviations: Pfr.f = prefrontal facet; am = anterior process of maxilla; asm = ascending process of maxilla; tcm = transverse crest of maxilla.

### Parietal

Posterodorsally articulates to the supratemporal and exoccipital. Anterodorsally articulates with the frontals. Posteromedially articulates to the prootic. Anteroventrally articulates with the basioccipital and basisphenoid.

#### Dorsal

Anterior side exhibits two transverse processes, whose lateral extensions make up the widest part of the bone. Transverse processes flare posterolaterally after the anterodistal terminus. Posterodistal terminus forms the ventrally oriented postorbital processes. From the posterodistal terminus the transverse processes curve anteromedially back toward the body of the bone. After meeting the body there is no curvature and the posterior end of the bone tapers to a shallow notch. The shallow notch between two rounded, short processes forms the posterior extension of the parietal. Ventral to the transverse processes lateral extensions are visible on both sides (Fig 11).

#### Ventral

The center of the parietal is moderately concave for ~8 mm toward the posterior, with ventral processes on each lateral edge. Postorbital processes form a ridge that curves posteromedially forming the lateral extensions and curving back toward the body anteriorly to meet the central processes. Posterior to the ventral processes the dorsal attachment to the body’s base exhibits a wavy, wrinkled appearance. Once meeting the body, the dorsal attachment flushes evenly with the body. The shallow notch exhibits borders between processes and the processes exhibit short projections at the anterior curvature.

#### Ontogeny

In ventral view the postorbital process is ventrally oriented in AR12, in AR2 and AR1 it is posteroventrally oriented (Figs 12 and 13). The anterior margin comes to a sharp point in AR2 while it is more of a smooth outward curve in AR12. Ventral processes are well developed in AR12, but in AR2 the anterior of the processes is more anteriorly pointed and the medial margin is less smooth. A flat, straight area is present on AR1 where the ventral processes would be located. Anteriorly this area is softly rounded and curved anteriorly to meet the frontal facets. Dorsoventral attachment to the body is straight in AR1, mostly straight in AR2, and in AR12, there are undulations. Posterior processes extend most in AR12 while in AR1 the area is flat and straight. In AR1 the body is open, no structures present but fossa on either side posteriorly. Transverse processes are not visible when viewing in ventral view. Lateral extensions of the posterior of AR1 are shorter and the medial margin extends first anteromedially then anteriorly, while in AR2 the medial margin curves anterolaterally to meet the ventral processes. In AR2 and AR12 the lateral extensions extend posterolaterally while in AR1 they extend posteriorly. Anterior transverse processes are much wider/thinker in AR1 than the others. In dorsal view the posterior is flat in AR1, shallow with a thin notch on AR2 and shallow with a wide notch in AR12. Posterior processes exhibit a rounded margin in AR12; in AR2 undulations are present. AR2 has many undulations on posterior border while others exhibit a flatter border. Fossa of AR1 are more visible in dorsal view, transverse processes are thinner and longer in AR12 and AR2, and in AR1 they are short and thick. Lateral extensions are not easily visible in dorsal view of AR1, but they are posteriorly oriented and close to the body.

##### Summary of Ontogeny

- Ventral processes begin as a flat edge in AR1, appear as anteriorly oriented, sharply pointed processes in AR2, and anterolaterally oriented, round-tipped processes in AR12.
- Dorsoventral attachment to body from ventral processes begins straight and develops undulations with age.
- Body of parietal in AR1 has not fully ossified, exhibiting two openings.
- Medial margin of lateral processes shift from curving anteromedially to meet the ventral processes in AR1 to anterolaterally in AR2 to mostly straight anteriorly in AR12.
- Transverse processes elongate with age and begin thick in AR1, thin in AR2, and thicken in AR12.
- In dorsal view the posterior margin is flat in AR1, shallow with a thin notch in AR2, and shallow with a wide notch in AR12.
- Processes of posterior margin begin flat, develop undulations, and round with age.

**Fig. 11.**
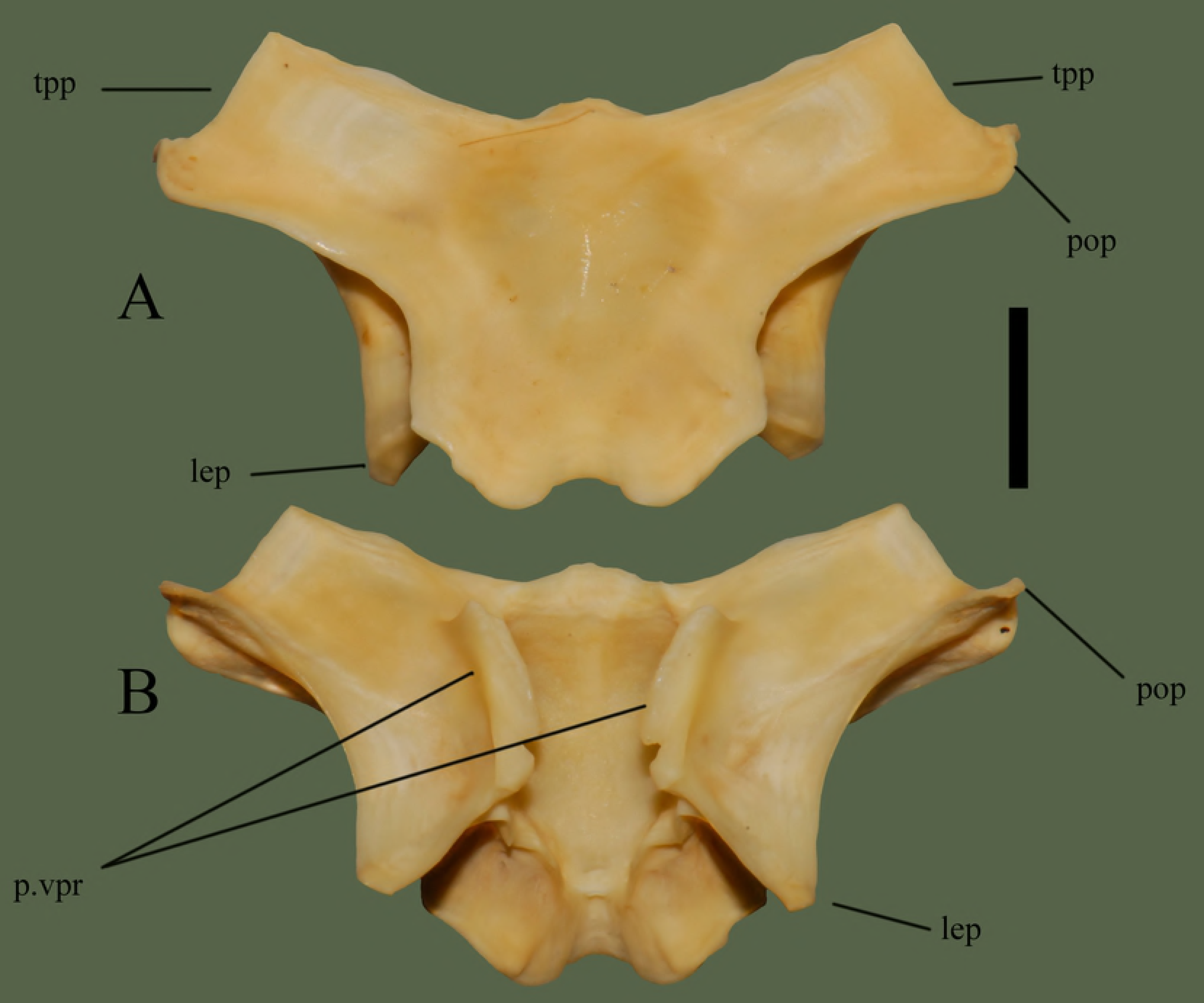
Parietal (ETVP 3295) in dorsal (A) and ventral (B). Anterior is to the top. Scale bar = 5 mm. Abbreviations: p.vpr = parietal ventral process; lep = lateral extension of parietal; pop = postorbital process of parietal; tpp = transverse process of parietal.

**Fig. 12.**
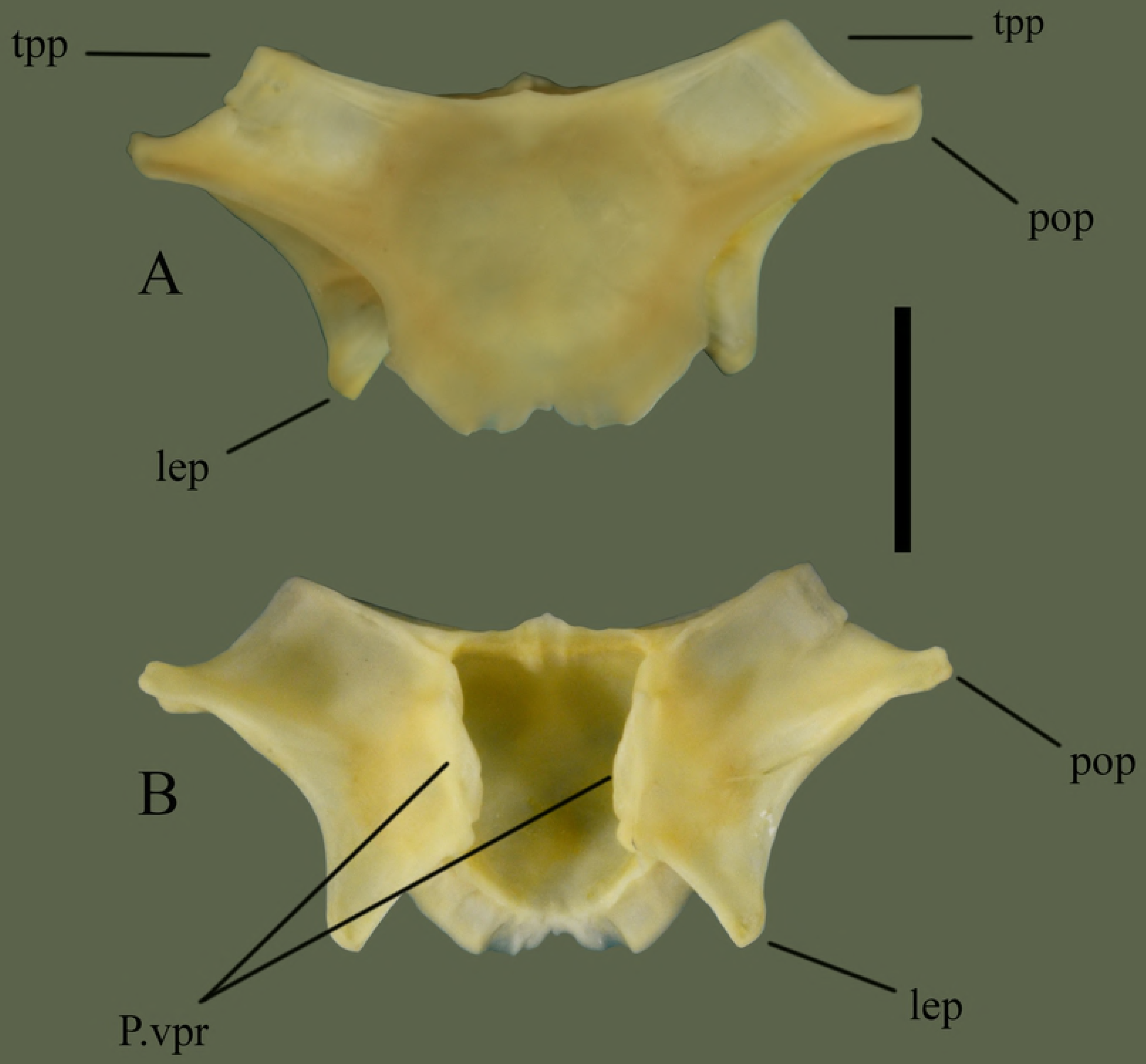
Parietal (ETVP 3293) in dorsal (A) and ventral (B). Anterior is to the top. Scale bar = 5mm. Abbreviations: P.vpr = parietal ventral process; lep = lateral extension of parietal; pop = postorbital process of parietal; tpp = transverse process of parietal.

**Fig. 13.**
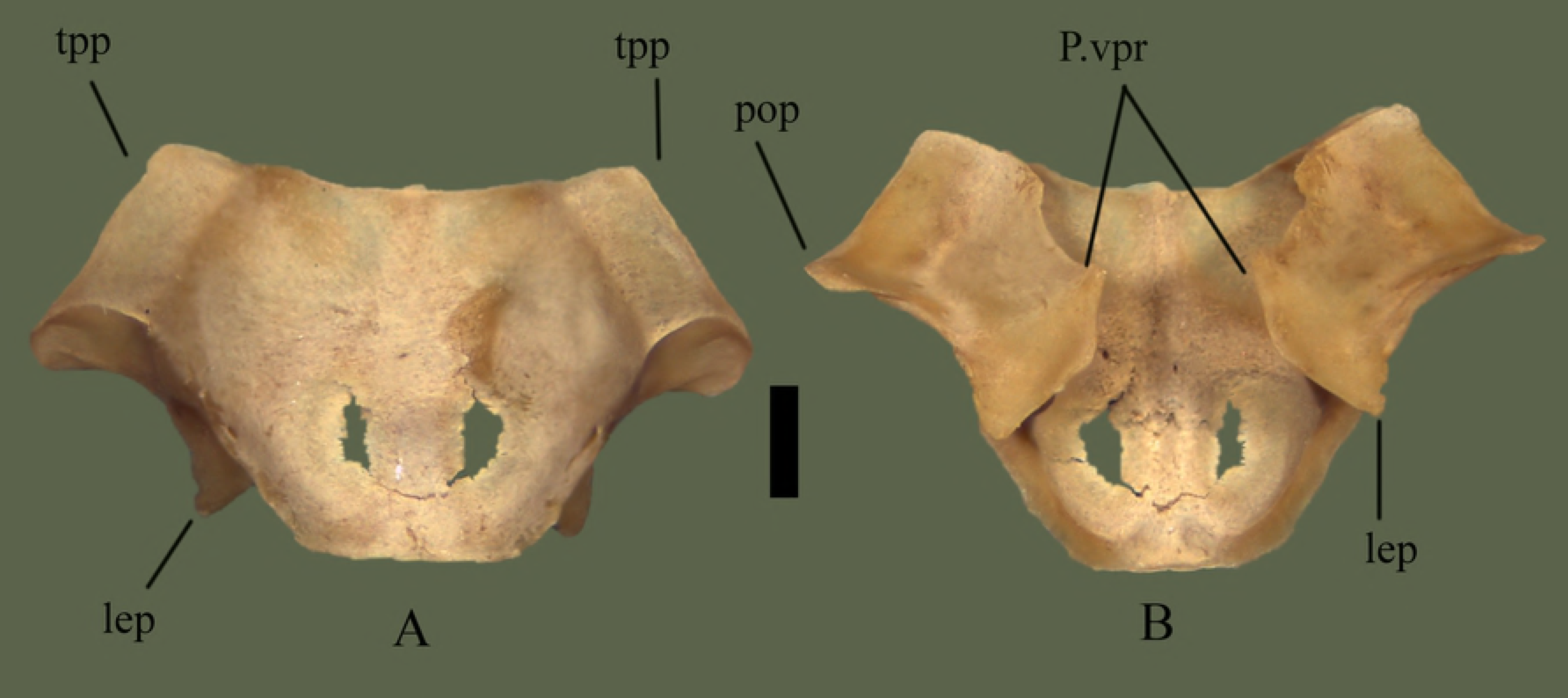
Parietal (ETVP 3306) in dorsal (A) and ventral (B). Anterior is to the top. Scale bar = 1mm. Abbreviations: p.vpr = parietal ventral process; pop = postorbital process of parietal; lep = lateral extension of parietal; tpp = transverse process of parietal.

### Prefrontal

Articulates anteroventrally to the maxilla; the anterior facing maxillary facet of the prefrontal sits in the cupped articulation surface at the dorsal end of the ascending process of the maxilla. The prefrontal articulates posterodorsally to the frontal; the frontal facet on the posterodorsal surface of the prefrontal fits into, and slightly ventral to, the prefrontal facet on the anterior of the frontal.

#### Dorsal

Medial side is elevated, extending posteriorly over the prefrontal edge. An elevated knob is present slightly lateral to the center. Lateral to the knob the edge is a rounded and ventrally oriented lateral process. A fossa is present near the center of the bone. Anterior section is somewhat flat laterally, and angled medially, with the flat area containing the maxilla facet. Posterior edge is curvy, containing the frontal facet (Fig 14).

#### Ventral

Lateral side is concave with a short ridge, formed by the lateral process, outlining the lateral edge. Medially the surface curves dorsally causing the medial side to be slightly elevated. Medial side is ventral to the lateral edge, though it is still somewhat concave in the center. Anterior and posterior edges are elevated forming short, rounded processes. Posterior process is slightly more elevated than the anterior. Another process is visible medial to the posterior process. The flat/depressed area is the articulating surface for the maxilla. Posterior edge is curvy with four projections. Medial to the cup on the lateral side is a short projection. Dorsal to the posterior process are two processes the medial one being more prominent and the lateral one being more flat.

#### Ontogeny

In dorsal view the lateral side of AR1 extends further out than others. In AR2 the lateral process is close to the knob, as in AR12 (Figs 15 and 16). In AR12 the lateral margin is rounded posteriorly then angles medially at anterior to meet maxillary facet. In AR2 the rounded posterior makes up less of the lateral margin, with a straight section anteriorly that then angles medially to curve into maxillary facet. In AR1 the lateral margin is rounded but thin posteriorly, then curves anteromedially, gently first, then more sharply to the maxillary facet. Maxillary facet is straight in AR12, in AR2 and AR1 it curves outward anteriorly. Anterior medial process is rounded, thin, and long in AR1. In AR2 it is shorter, thick and rounded. In AR12 it is more square-shaped, short, thick, and oriented more medially. Posterior medial process is elongate, flat tipped, and curved more anteriorly than posteriorly in AR1. In AR2 it is elongate, rounded, and posterior side curves in anteriorly to itself. In AR12 it is short, the tip is square shaped with rounded corners, and the sides curved gently to meet the body. Dorsal knob is thinnest in AR1 and extends posteriorly more than others. Posterior of the knob is oriented posteromedially with a squared margin in AR1. Anterior process of the knob is anteroposteriorly long and tallest in AR1. A fossa is present anteromedially to anterior process of the knob, the area surrounding the fossa in concave in AR1. In AR2 the knob is mostly rounded and extends posteriorly, more than AR12 but less than AR1. Anterior process is much shorter than AR1 but still taller than AR12. In AR12 the anterior process is shortest. Posterior margin of knob has a rounded protrusion that curves anterolaterally to meet body. The knob extends posteriorly the least in AR12. Posterior margin is straight with little variations in AR1. In AR2 undulations are more exaggerated while in AR12 they are more gently curved. In ventral view the lateral ridge is most sharply curved in AR12, most gently in AR1. Lateral side is elongate and thin in AR1, thick in AR2, and thickest in AR12. Concave area of lateral side is smooth in AR12, rough and deep in AR2 and in AR1 appears as a lowered area rather than fully concave. Anterior process of medial side extends furthest in AR1, is shorter and wider in AR2, and in AR12 is a curved corner with a short rounded protrusion. Posterior medial process is long in AR1, tip comes to a rounded point, and a ridge is present ventrally that curves out posteriorly. In AR2 posterior medial process is very rounded, extends from the body, and ridge is present ventrally but does not extend posteriorly. In AR12 the posterior medial process is short, wide, and anterior side extends farthest. The ventral ridge has rounded and extended posteriorly to form another process in AR12. Posterior lateral process is rounded and extends from the body in AR1. In AR2 it is rounded, wider, and meets the lateral ridge ventrally. In AR12 is it much thinner, meets ridge ventrally, and from the posterior it is angled medially to meet the body. The thicker, rounded area of the lateral ridge is present in AR1 as is the anterior knob. In AR2 the ridge is much thicker than AR1 and has curved in on itself. In AR12 the ridge is thickest and extends most anteroposteriorly.

##### Summary of Ontogeny

- Lateral margin begins thin, extending more laterally, then thickens and shrinks closer to the body of the bone with age.
- Maxillary facet begins anteriorly curved and becomes straight/flat by AR12.
- Posterior medial process rounds and recedes close to the body with age.
- Anterior process of dorsal knob shortens with age.
- Dorsal knob recedes closer to body anteriorly with age.
- Lateral side thickens with age.
- Lateral ridge curves gently in AR1 while curving sharply in AR12.
- Anterior process of medial side shortens and recedes into body of bone until it becomes a rounded corner with a short, rounded, protrusion.
- Posterior medial process shortens/recedes into the body and widens with age.
- Ridge on posterior medial process begins oriented posterolaterally, in AR2 extends ventrally, and in AR12 is present as a posterior facing rounded process.
- Posterior lateral process widens and recedes closer to the body with age.
- Lateral ridge thickens and elongates anteroposteriorly with age.

**Fig. 14.**
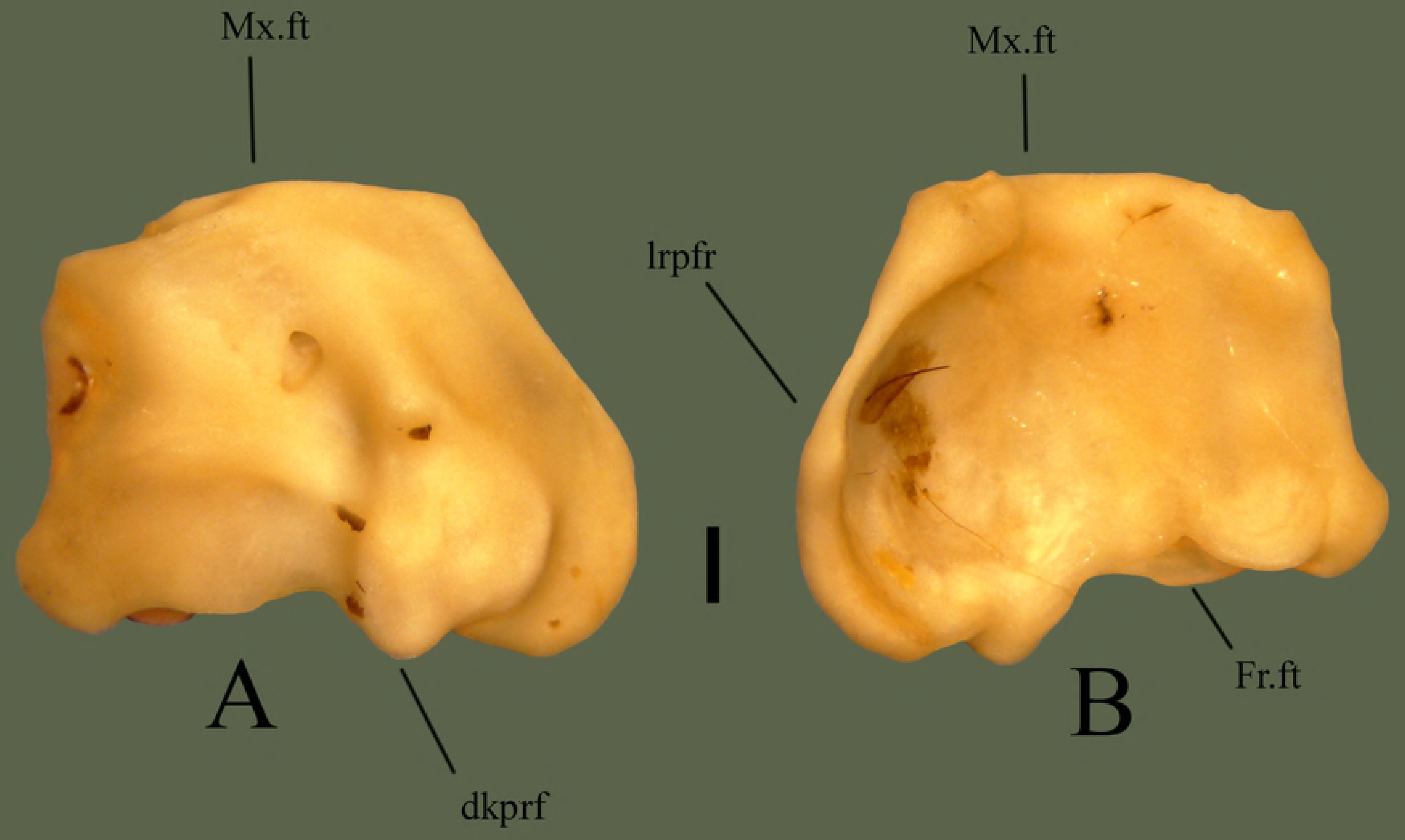
Right prefrontal (ETVP 3295) in dorsal (A) and ventral (B) view. Anterior is to the top. Scale bar = 1mm. Abbreviations: Mx.ft = maxillary facet; Fr.ft = frontal facet; dkprf = dorsal knob of prefrontal; lrpfr = lateral ridge of prefrontal.

**Fig. 15.**
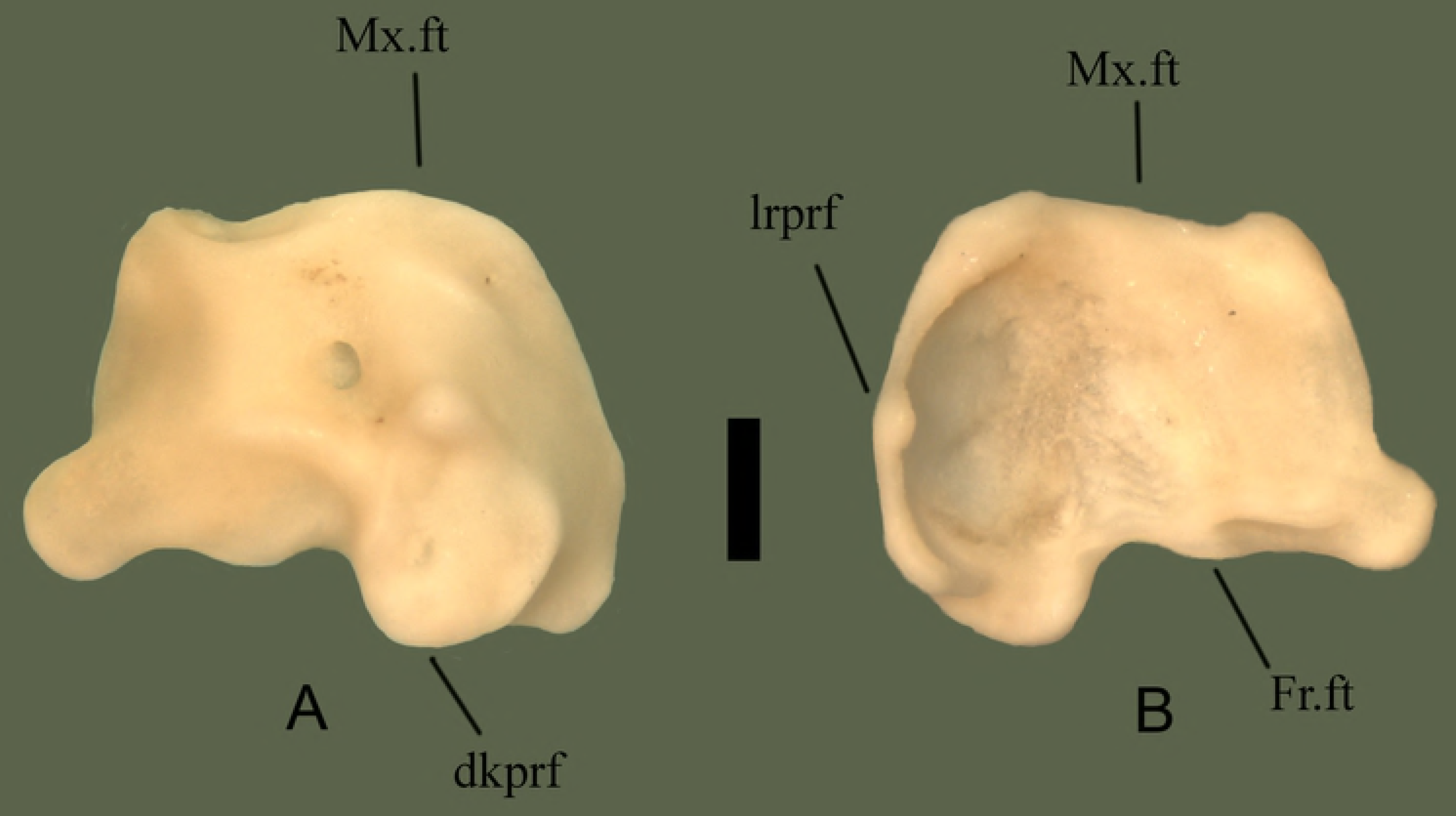
Right prefrontal (ETVP 3293) in dorsal (A) and ventral (B) view. Anterior is to the top. Scale bar = 1mm. Abbreviations: Mx.ft = maxillary facet; Fr.ft = frontal facet; dkprf = dorsal knob of prefrontal; lrprf = lateral ridge of prefrontal.

**Fig. 16.**
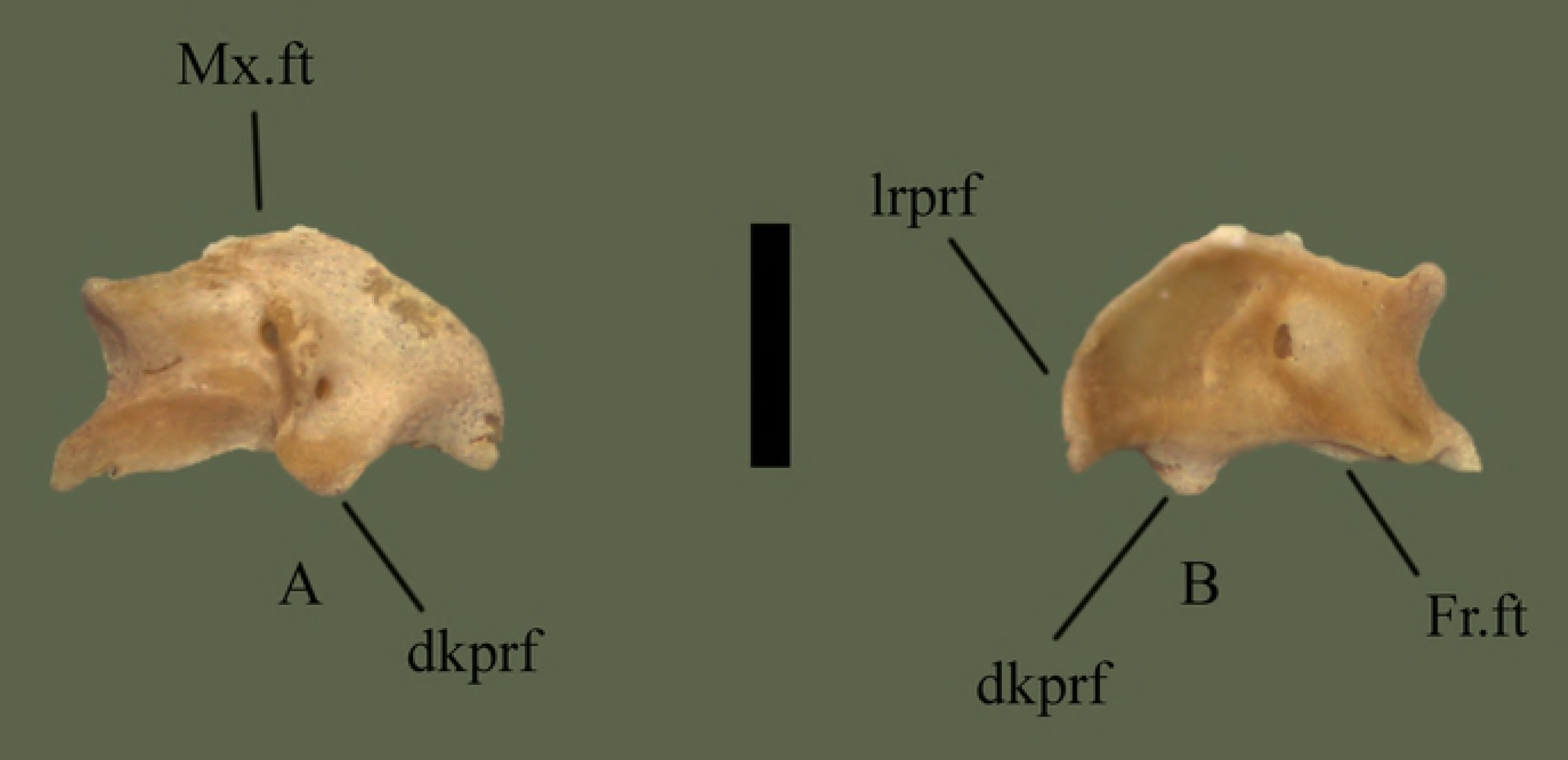
Right prefrontal (ETVP 3306) in dorsal (A) and ventral (B) view. Anterior is to the top. Scale bar = 1mm. Abbreviations: Mx.ft = maxillary facet; Fr.ft = frontal facet; dkprf = dorsal knob of prefrontal; lrprf = lateral ridge of prefrontal.

### Premaxilla

The premaxilla articulates posterodorsally to the septomaxilla; the anterior tips of the septomaxilla sit on either side of the ascending process of the premaxilla. The premaxilla articulates posteroventrally to the vomers; the posteriorly facing vomerine process of the premaxilla abuts the anterior tips of the vomers.

#### Dorsal

An ascending process is present on the dorsal side, is ~1 mm tall, and does not meet the nasals. Premaxilla exhibits well developed transverse processes that extend ~3 mm from the center of the anterior edge. The vomerine process makes up the posterior end, extending out ~1mm and tapering posteriorly. A fossa is present on each distal side of the ascending process with one extra fossa on the left side (Fig 17).

#### Anterior

Ascending process is dorsally oriented, rounded, and thick. Ventral side of the transverse process widens medially, reaching maximum a width of ~1.5 mm from the distal tip before curving inward dorsally.

#### Ontogeny

Vomerine notch deepens and vomerine processes extend with age. Fossa present on either side of the ascending process; on AR1 fossa are close to parallel, in AR2 the left one is smaller and positioned more medial, and in AR12 the right fossa is barely visible and positioned anteriorly. Anterior edge of AR1 is a single, gentle curve; AR2 has formed the double undulated shape that is seen in AR12. On AR1 a ventral fossa is present, with AR2 appearing concave in the same area, and no evidence of either on AR12.

##### Summary of Ontogeny

- Vomerine notch deepens with age.
- Vomerine processes extend with age.
- Ventral process disappears with age.
- Anterior margin starts as a single gentle curve but by AR2 exhibits the double undulated shape also seen in AR12.

**Fig. 17.**
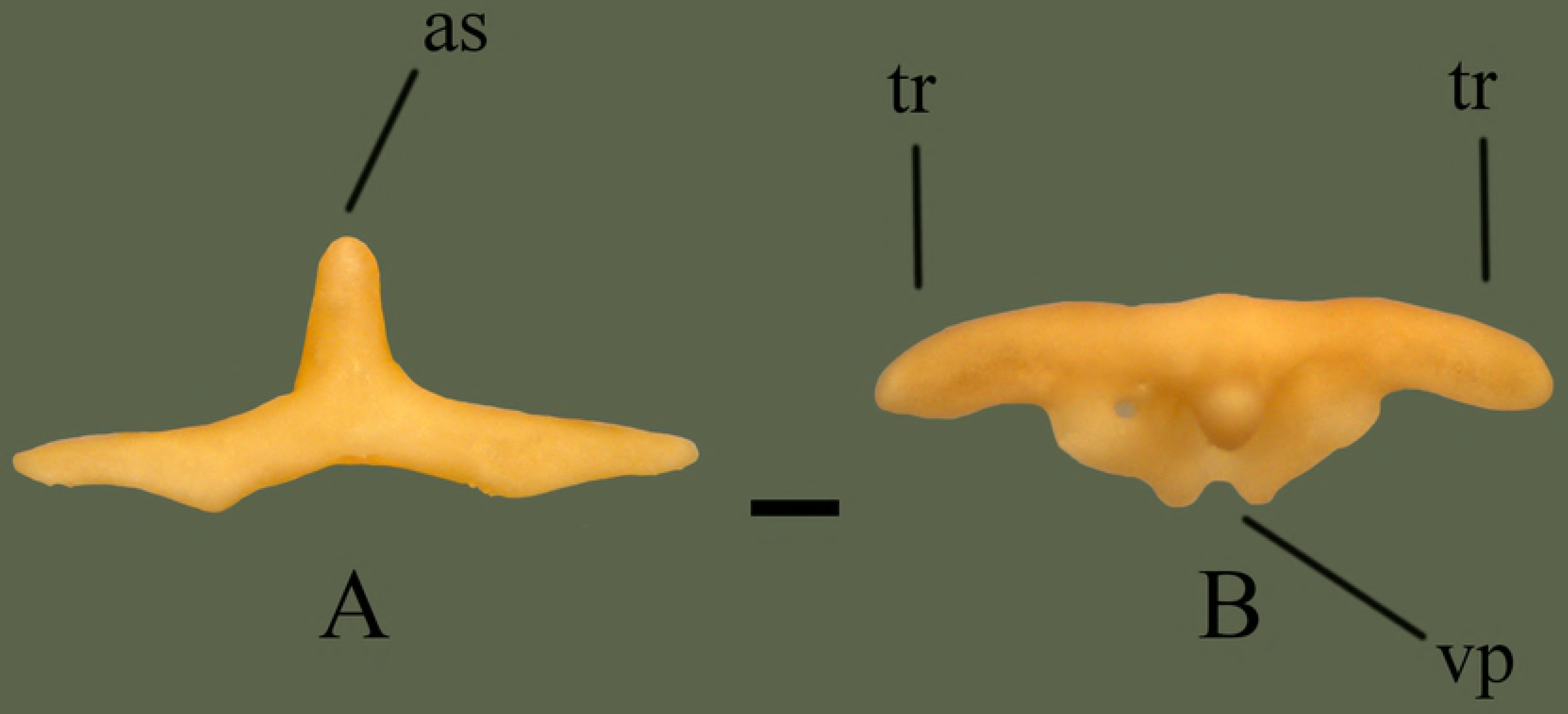
Premaxilla (ETVP 3295) in anterior (A) and dorsal (B) view. Anterior is to the top. Scale bar = 1mm. Abbreviations: as = ascending process of premaxilla; tr = transverse process of premaxilla; vp = vomerine process of premaxilla.

### Dentary

Posteriorly the dentary articulates with the compound bone, the angular, and the splenial. Anterior of the splenial fits into the medial groove of the dentary, with the dorsal process of the splenial connecting to the ventral half of the medial process of the dentary. Anterodorsal process of the angular connects to the dorsal half of the medial process of the dentary, so that the medial process connects with the angular on it’s dorsal half and the splenial on it’s ventral half. Ventral process of the dentary also borders the anteroventral edge of the angular. The anterior of the compound bone fits between the ventral and dorsal processes of the dentary, enclosing it’s lateral groove.

#### Medial

Anterior end is curved medially while the body remains uniform. Teeth are aligned to the lateral side. A narrow groove is present on the medial side, open on the posterior end and present as an indented line on anterior end. Dorsal and ventral edges of the groove curve in on themselves dorsoventrally. The dorsal edge projects out medially producing a curved process with a back and forth curved posterior end. This dorsomedial process articulates with the dorsomedial process of the compound bone. As in lateral view a dorsal and ventral process are produced by the bifurcating of the posterior end of the dentary (Fig 18).

#### Lateral

The right dentary exhibits ~13 teeth that span the entire length, with one at the very posterior. Anterior teeth are the largest, with a decrease in size posteriorly. Posteriorly the dentary bifurcates into thin dorsal and ventral processes. The center of the bone bears a deep groove with the posterior half being open and the anterior half having bone visible from the lateral side. Anterior end of the compound bone articulates with the groove. Anterior to the groove is a deep fossa.

#### Ontogeny

In lateral view the anterior tip is somewhat ventrally angled on AR2. Anterior tip is curved less in AR1 and curved the least in AR12. Anterolateral fossa is anteroposteriorly long on AR1, short and shallow on AR2, and the most shallow on AR12. Ventral process on the posterior of AR1 is curved dorsally. Tooth attachment areas are visible on AR1, perhaps not yet sockets. In medial view the ventral and dorsal folds on the anterior of AR1 do not meet. The folds begin to meet in AR2, and completely meet in AR12. Dorsomedial process of AR2 exhibits two short, undeveloped processes. In AR1 the two processes are visible, close together, and taller than in AR2. In AR12 they are developed, rounded, and protrude more than in AR2 and are thicker than both AR1 and AR2.

##### Summary of Ontogeny

- Anterior lateral fossa shortens anteroposteriorly, becomes circular in shape, and shallows with age.
- Ventral posterior process is angled dorsally in AR1, extends posteriorly in others.
- Dorsal and ventral folds on medial side get closer with age, eventually meeting/joining in AR12.
- Dorsomedial process of dentary begins thin and elongate, thickens and shortens in AR2, then elongates and thickens in AR12.

**Fig. 18.**
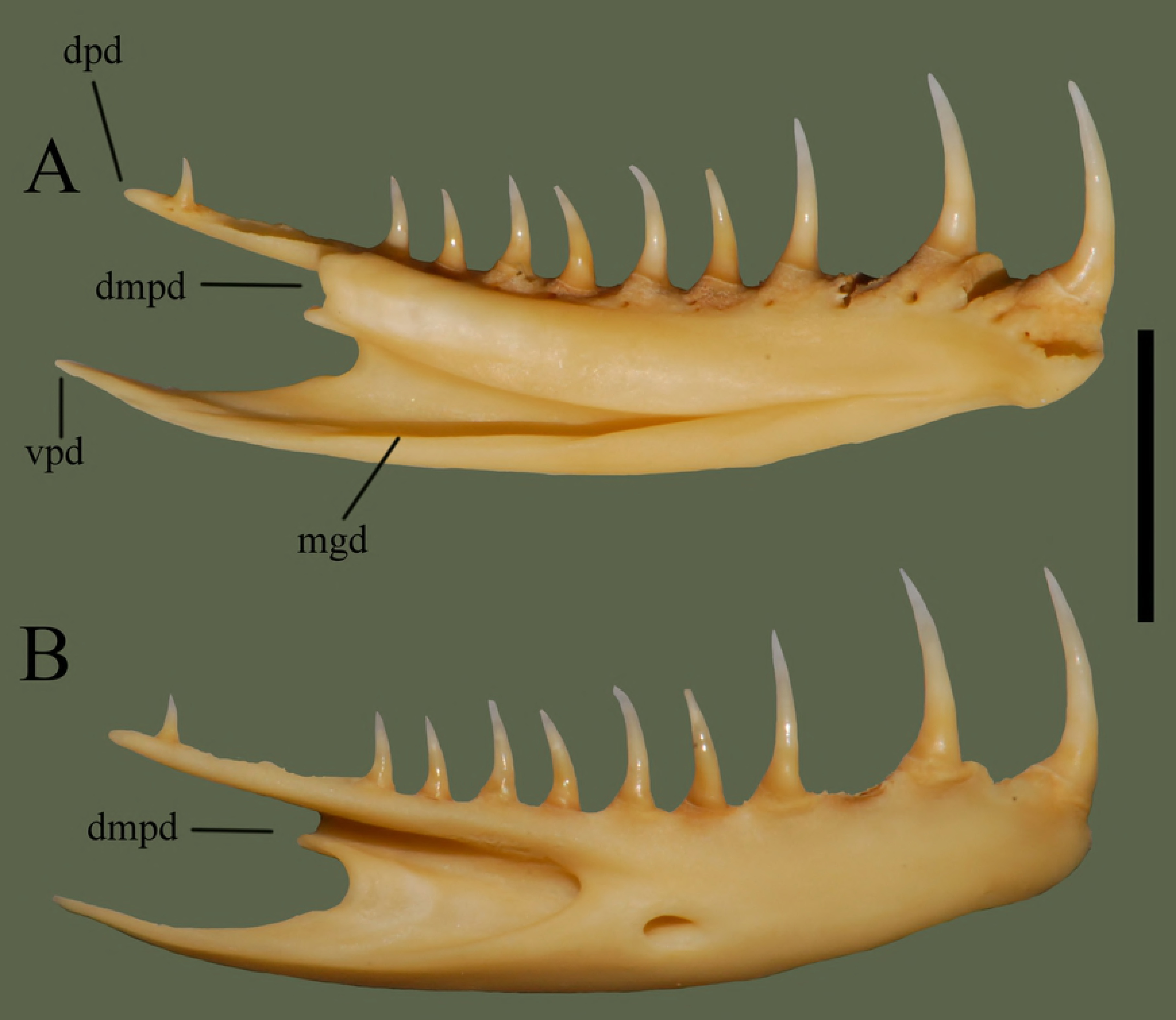
Right dentary (ETVP 3295) in medial (A) and lateral (B) view. Anterior is to the right. Scale bar = 5 mm. Abbreviations: dpd = dorsal process of dentary; dmpd = dorsomedial process of dentary; mgd = medial groove of dentary; vpd = ventral process of dentary.

### Angular

The anteroventral edge of the angular borders the ventral process of the dentary. The anterodorsal process of the angular articulates with the dorsal half of the medial process of the dentary. The narrow posterior of the angular fits into the medial groove of the compound bone. Anterior of the angular connects to the posterior edge of the splenial.

#### Medial

Anteriorly there is a dorsally oriented process, representing the anterior-most end of the bone. The process exhibits a flat edge and curves back posteroventrally, causing a widening of the ventral side. Approximately 2 mm from the anterior edge the curve terminates at a rounded process that is oriented ventrally. Anterior and posterior edges of the rounded process extend past the base of the process, forming a “pinched” appearance at the base. Directly dorsal to the rounded process is a small, triangular process. Posterior to the rounded process the bone tapers dorsally, thinning the posterior section. Posterior end consists of a rounded point (Fig 19).

#### Lateral

Ventral edge appears thicker and elevated. The elevated ridge runs posteriorly from the rounded process, terminating as the rounded posterior end. Bone dorsal to the ridge is gently concave with some elevation occurring at the most dorsal section of the dorsal process. The bone also appears thinner in the same areas, with the exception of the most dorsal section of the dorsal process. The rounded process does not appear as protruding or rounded in lateral view. Dorsally the ridge terminates with a slightly rounded tip, flaring ventrally, widening the anterior of the ridge. Approximately 1 mm from the anterior-most of the rounded process the curving of the anterior of the ridge end in a sharp point and curves anterodorsally toward the body of the bone. Posterior of the ventral side curves lightly with no other significant features. Dorsal to the rounded process and ventral to the dorsal process the triangular process is more visible in lateral view. Dorsal and ventral edges of the triangular process curved in their respective directions to meet the body of the bone.

#### Ontogeny

Dorsal process of AR1’s right angular has broken off (not ontogenetic). Posterior tip is sharply pointed on AR1 and AR2; on AR12 it is rounded. Dorsal margin exhibits most curvature in AR12 with AR2 being less curvy and AR1 the least curvy. Dorsal process is thickest in AR12; in AR2 it is much thinner and less dorsally oriented. While the dorsal process of the right angular of AR1 has been broken, the left angular shows a thin dorsal process, like in AR2, that is less dorsally oriented than in AR12. Tip of the dorsal process is rounded in AR1, square in AR12, and intermediate in AR2. In lateral view of AR2 the ventral process is thinner anteroposteriorly than AR12. Groove of the ventral process is formed in AR2 and the termination point is more anterior than in AR12. Ventral process of AR1 is elongate and exhibits a groove; dorsal margin of the groove is long, unlike in AR2 and AR12. Ventral margin gently curves out ventrally in AR2. In AR1 ventral margin is straight. On AR1 the lateral side is more concave than in AR12 and AR2, with AR12 being the least concave. Anterior rounded process is “pinched off” in AR12, inward curving is present on AR2, posterior to the process, but does not exhibit the “pinched” shape. In AR1 the ventral margin of the anterior rounded process is straight with no “pinching”. Curvature from dorsal to ventral process is square in AR1, deeply curved in AR2 and gently curved in AR12.

##### Summary of Ontogeny

- Dorsal margin increases in curviness with age.
- Dorsal process of angular thickens and protrudes more dorsally with age.
- Posterior margin of dorsal process becomes more square-shaped with age.
- Lateral side of angular becomes less concave/more flat with age.

**Fig. 19.**
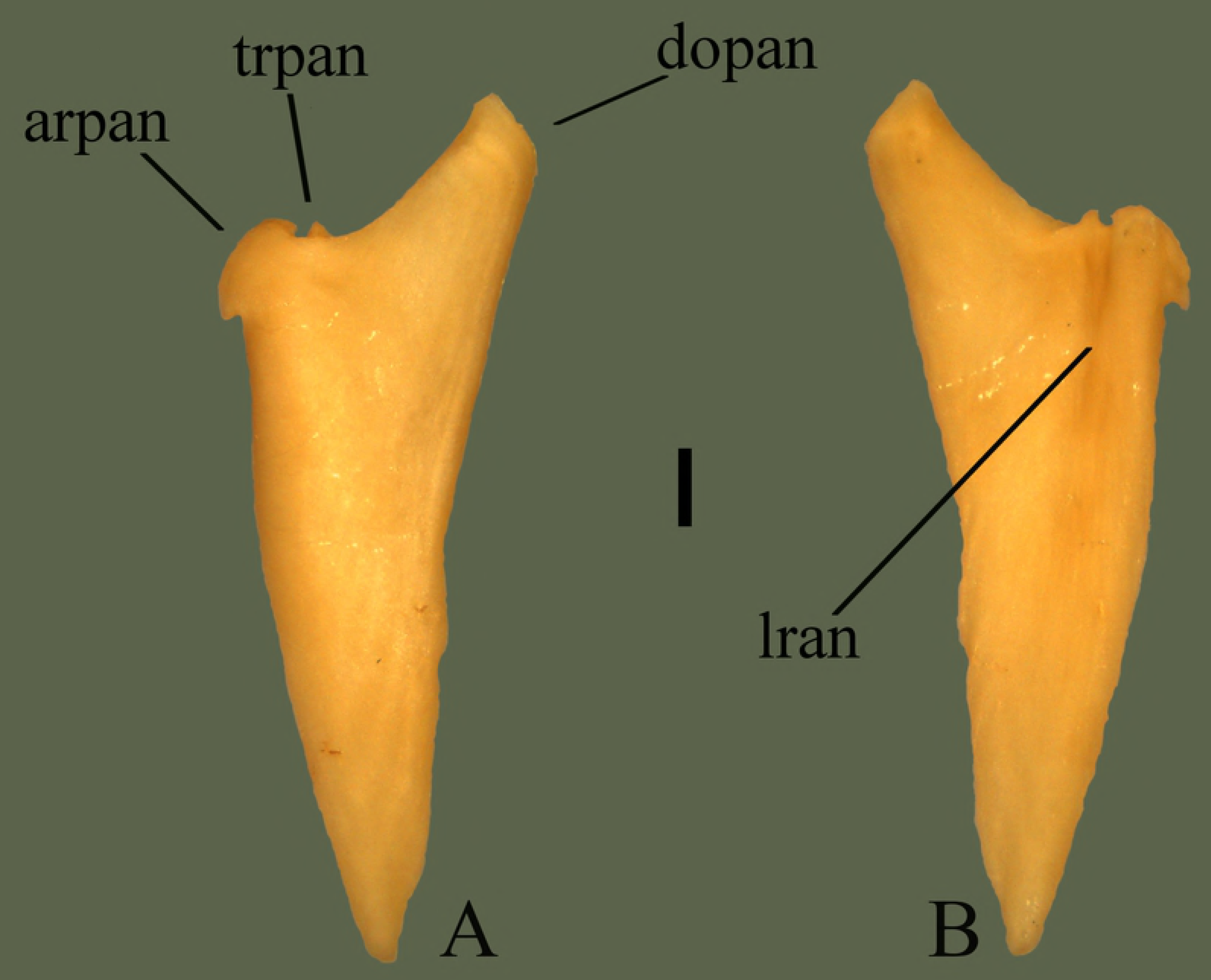
Right angular (ETVP 3295) in medial (A) and lateral (B) view. Anterior is to the top. Scale bar = 1 mm. Abbreviations: arpan = anterior rounded process of angular; dopan = dorsally oriented process of angular; lran = lateral ridge of angular; trpan = triangular process of angular.

### Splenial

The posterior edge of the splenial articulates to the anterior edge of the angular. The tapering anterior of the splenial fits into the medial groove of the dentary, leaving only the anterior mandibular foramen open.

#### Medial

Anterior end comes to a point, forming a bone that is anteriorly tapering with an expanded posterior end. From the anterior point the bone gradually widens posteriorly. Approximately 3 mm from anterior point the widening ends where the dorsal side comes to a point before curving inward ventrally, toward the body. The curve extends ~2.5 mm posteriorly, before then curving outward dorsally. An anteriorly oriented process is formed by the dorsal termination of the curve. Dorsal side of the process bears a small ridge, extending posteriorly and ending at the articulating end. A depression is observed on the dorsal edge of the dorsal process. The posterior end exhibits a curved process to articulate with the angular. Aside from the process, the rest of the posterior end has slight curvatures for articulation, but is mostly uniform. Ventral border of the bone is mostly straight, with slight curvature (Fig 20).

#### Lateral

A spine runs from the anterior-most tip, reduces in height ~3/4 way down toward posterior, and terminates at the center of the posterior notch. A fossa is present on the dorsal side of the spine. The posterior end exhibits more relief than in medial view, with the medial side of the posterior process extending past the posterior notch.

#### Ontogeny

Anterior tip is sharply pointed in AR12 and AR1 with AR2 being rounded, though this could be due to breakage. Anterior dorsal margin is straight and flat in AR12; AR2 shows little curvature and AR1 shows the most curvature; curving outward dorsally. Dorsal incisure is anteroposteriorly oriented in AR12, in AR2 there is some ventral curvature, and in AR1 there is a posteroventral curvature. Both AR1 and AR3 are curved on the ventral margin while AR12 is mostly straight. Posterior end of AR1 exhibits four undulations, ventral process is not yet extended. A fossa is present dorsal to the ventral process, not seen on AR12 or AR2. In medial view the lateral ridge is visible posteriorly in AR1 and AR2l the ridge does not fully extend anteriorly. In lateral view of the posterior the ventral process is present on AR2, but not fully extended posteriorly. Tip of the anterodorsal process is sharply pointed in AR1 and AR2, rounded in AR12. A short ridge is present on the dorsal edge of the dorsal process of AR1. The short ridge on the posterodorsal process of AR2 is slightly more ventral than in AR1. On AR12 the ridge is long and positioned most ventral. The lateral ridge is closest to the ventral margin in AR1. In AR2 the ridge is positioned more dorsally and in AR12 the ridge terminates and the anterior tip. The section of bone ventral to the lateral ridge appears to thin with age. The posterior notch on AR12 is curved anteriorly and the medial side of the notch extends posteriorly past the lateral side. The notch on AR2 has a “V” shaped indentation on the more dorsal side and the medial side extends past the lateral side of the notch. A notch has not yet formed on AR1, the lateral ridge extends past the posterior edge, and the ventral margin has many undulations. Margin of the incisure in AR2 is the most undulating.

##### Summary of Ontogeny

- Anterior dorsal margin straightens/flattens with age.
- Dorsal incisure shifts from posteroventral curvature, to subtle ventral curvature, to anteroposteriorly oriented.
- Ventral process extends with age.

**Fig. 20.**
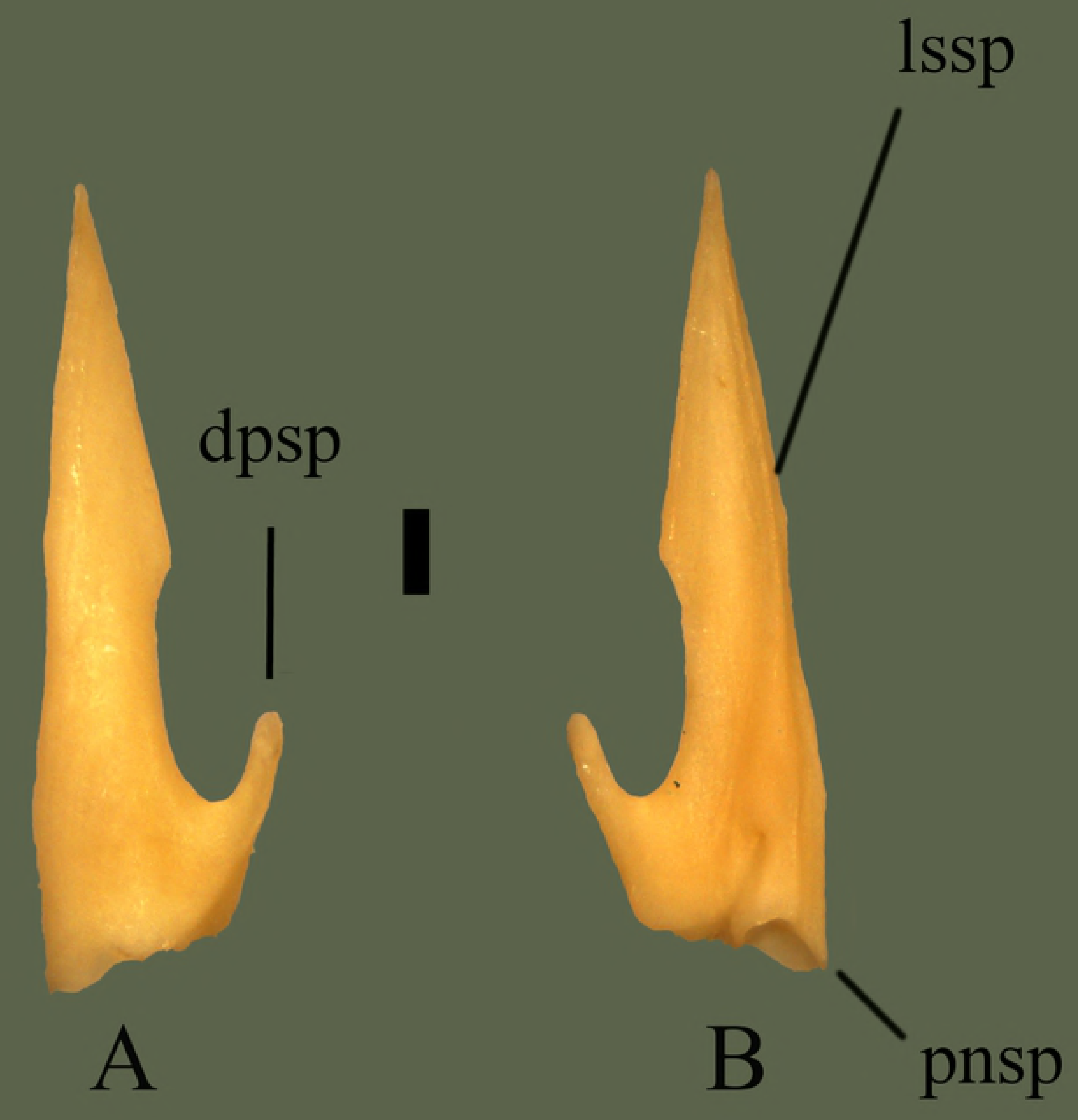
Right splenial (ETVP 3295) in medial (A) and lateral (B) view. Anterior is to the top. Scale bar = 1 mm. Abbreviations: dpsp = dorsal process of splenial; lssp = lateral spine of splenial; pnsp = posterior notch of splenial.

### Compound Bone

Anterior tip of the compound bone articulates to the dentary between the ventral and dorsal process, enclosing the lateral groove of the dentary. Medial groove of the compound bone receives the posteriorly tapering end of the angular. The articular of the compound bone interlocks with the mandibular condyle of the quadrate while the posterior of the pterygoid abuts the articular medially.

#### Medial

Anterior end comes to a point consisting of a lateral ridge and two medial ridges. Medial ridges curve in laterally, causing the junction to be concave. The concave area is mostly open, except a thin anterior process. Dorsal to the process is a relatively short dorsally elevated ridge. Apptoximately 13 mm from the tip of the anterior process the posterior begins to curve dorsally; widening the bone and marking the anterior of the prearticular crest. Prearticular crest curves out medially but does not extend past the width of the bone. Articular surface is more visible in medial view, demonstrating the texture and shape of the surface. Ventral border of the articular exhibits two fossa. Ventral to the anterior crest of the articular is a dorsomedially oriented process (Fig 21).

#### Lateral

Anterior end comes to a point, gradually widens posteriorly and maintains its width for ~2 mm before the bone begins to curve outward dorsally, forming the prearticular crest. Posterior to the prearticular crest the bone curves inward ventrally then outward dorsally, forming the anterior process of the articular. Articular surface is ~1 mm deep between two processes. Posterior to the articular is the ventromedially curved retroarticular process. The center of the prearticular crest exhibits a deep mandibular fossa. Anterior to the retroarticular process the ventral border curves out laterally, representing the surangular crest. Anterior to the termination of the surangular crest the bone remains uniform in width until the thinning and termination of the anterior end. Lateral side bears two fossa; Anteroventrally to the prearticular crest is a small fossa, anterior to that fossa is another smaller fossa.

#### Ontogeny

In AR1 the lateral edges of the prearticular crest are very rough. Mandibular fossa present in all Age Ranks. Fossa anterior to the mandibular fossa is present in all, a third fossa is present in lateral view of AR2. Third fossa is absent in AR1 and while the right compound bone of AR12 does not exhibit a third fossa, the left does. Anterior tip very sharply pointed in AR1. In AR2 and AR12 the anterior tip is rounded and edges are smooth. Medial view anterior end of AR1 very thin, bent, flimsy—well formed in AR2. Posterior fossa inferior to articular is large in AR1, perhaps more “open”. In AR2 it is not as open; instead it is narrow and deep. In AR12 the posterior fossa inferior to articular is the most deep and narrow. Anterior fossa of the posterior end is not present in AR2 and AR12. Retroarticular not yet fully formed in AR1, not curved, square-shaped, and dorsoventrally thinner than in AR12. Rounded process on retroarticular appears as a pointed process on AR1. Surangular crest is not yet formed on AR1.

##### Summary of Ontogeny

- Margin of the prearticular crest smooths with age.
- Surangular crest forms with age.
- Retroarticular shortens and thickens with age.
- Overall bone thickens with age.
- Posterior fossa of medial side narrows anteroposteriorly and deepens with age.

**Fig. 21.**
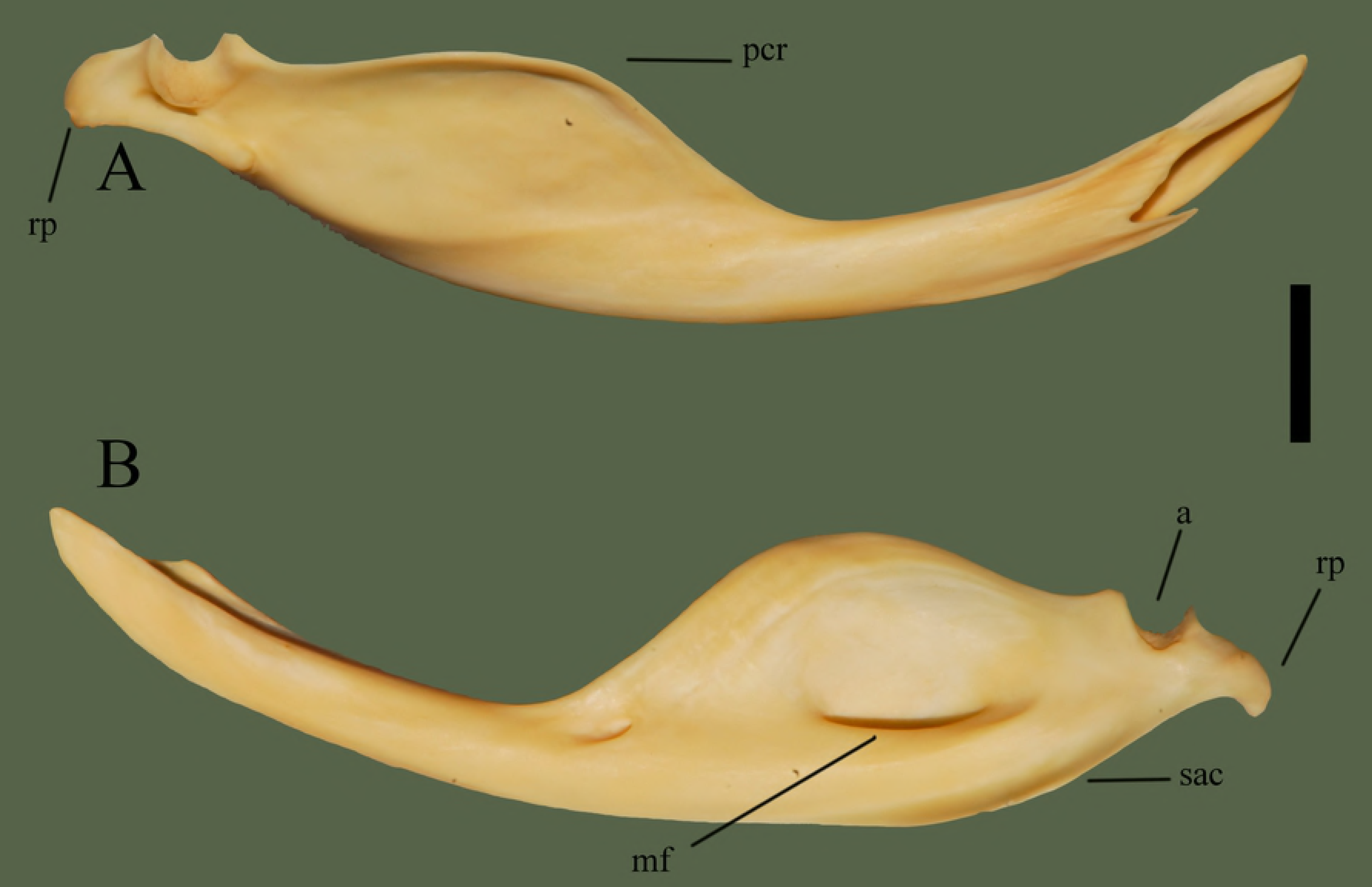
Right compound bone (ETVP 3295) in medial (A) and lateral (B) view. Anterior is to the right for A and to the left for B. Scale bar = 10 mm. Abbreviations: a = articular; mf = mandibular fossa; pcr = prearticular crest; rp = retroarticular process; sac = surangular crest of compound bone.

## Discussion

### Ontogeny

The goal for this study is to understand ontogenetic change and begin a long-term project of being able to identify isolated cranial elements in the fossil record by first providing a description of cranial elements and ontogenetic change in a single taxon.

Through ontogeny the bones that showed the most significant change were the prefrontal, maxilla, and parietal. The prefrontal shows an overall increase in robustness with age (Fig 22). The youngest specimens (AR1 and AR2) exhibited long, thin processes that would shorten and thicken with age (AR12). Edges with extreme undulations would become gentle undulations or thick processes. The maxillary facet curved outward anteriorly in the younger specimens, eventually becoming straight in AR12. This straightening could be due to the development of muscles and cartilage holding the prefrontal to the maxilla, allowing additional kinesis of the prefronto-maxillary joint. The maxilla exhibits a thickening of the ascending process with age, as well as a lengthening of the lateral process of the ascending process (Fig 23). Prefrontal facet shows significant change through ontogeny, in AR1 it is laterally rounded, medially straight, medially oriented, and dorso-ventrally elongate. In AR2 the facet becomes more rounded, and in AR12 it exhibits a rounded square shape. The change to a more rounded facet with age could also be due to the development of muscles, allowing a more kinetic articulation between the maxilla and prefrontal. Parietal exhibits a shift from a basic form to a more complex form (Fig 24). Postorbital processes are ventrally oriented in AR12 while in AR1 and AR2 they are posteroventrally oriented. Ventral processes are absent in AR1, yet in AR2 are present and their anterior is pointed and the medial margin is rough with many undulations. In AR12 the ventral processes exhibit a rounded anterior, and the medial margin is smooth. Ventral processes develop and “round out” with age. The lateral extensions show a slight shift in orientation, being posteriorly oriented in AR1 and shifting posterolaterally in AR2 and AR12. The transverse processes begin thick and short in AR1; with age they lengthen and subsequently thin out. The parietal of AR1 exhibits two fossa at it’s center; they exhibit rough edges, possibly indicating the continuing formation of the parietal. Aside from the mentioned characteristics, the overall trend appears to demonstrate small changes in proportions while the elements increase in size.

**Fig. 22.**
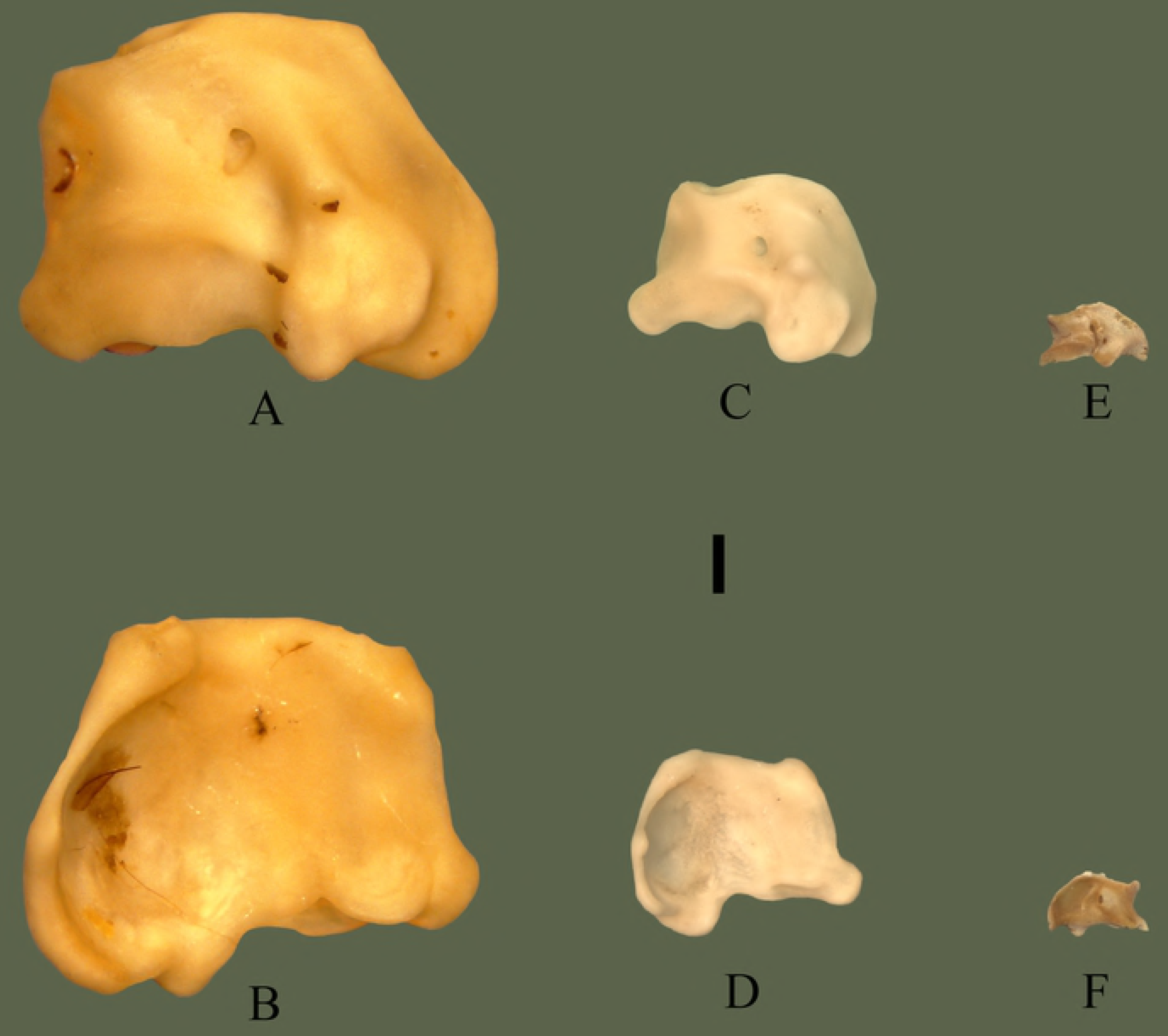
Prefrontal ontogeny. ETVP 3295 in dorsal (A) and ventral (B); ETVP 3293 in dorsal (C) and ventral (D); ETVP 3306 in dorsal (E) and ventral (F). Anterior is to the top of the page. Scale bar = 1 mm.

**Fig. 23.**
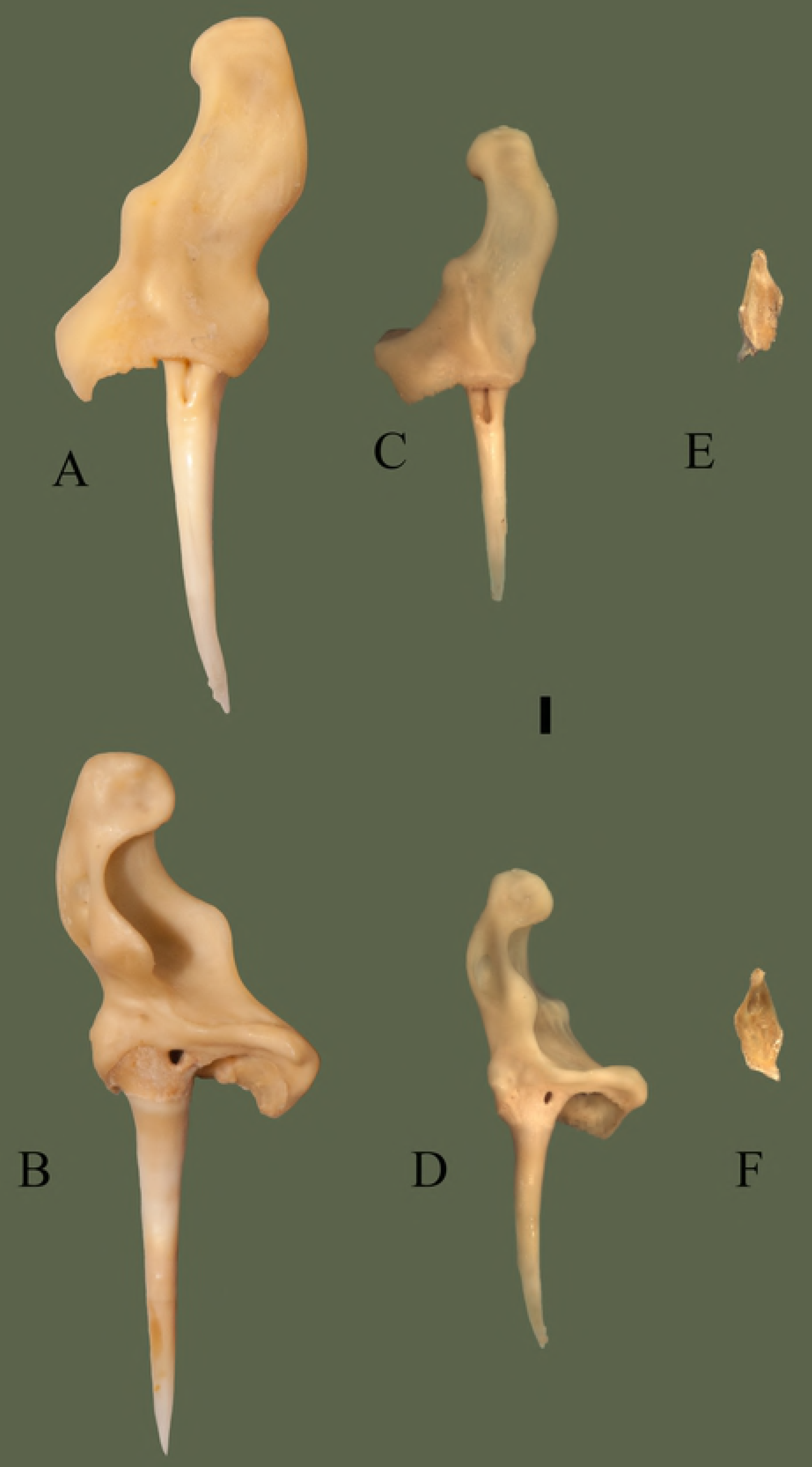
Maxilla ontogeny. ETVP 3295 in dorsal (A) and ventral (B); ETVP 3293 in dorsal (C) and ventral (D); ETVP 3306 in dorsal (E) and ventral (F). Dorsal is to the top of the page. Scale bar = 1 mm.

**Fig. 24.**
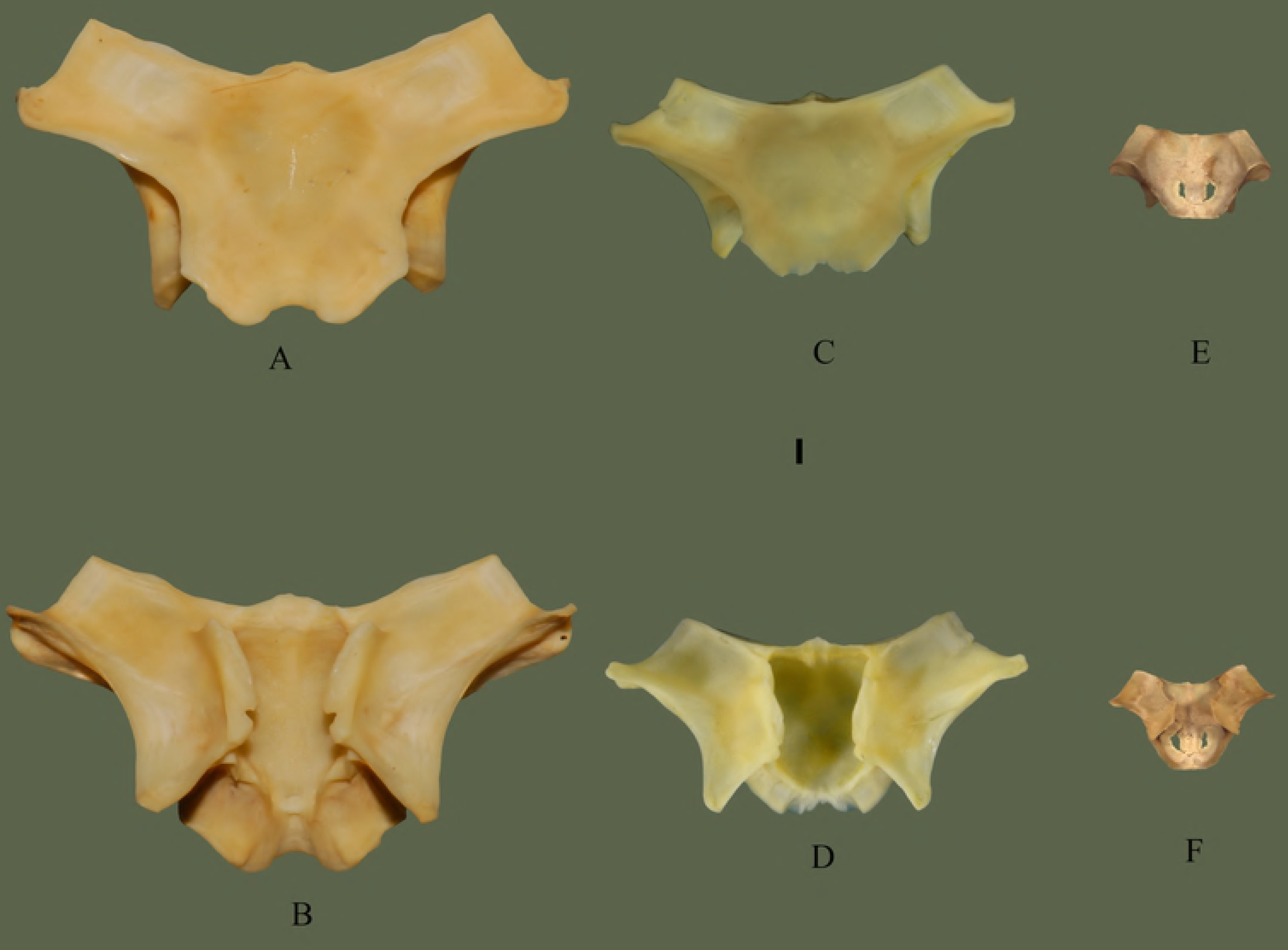
Parietal ontogeny. ETVP 3295 in dorsal (A) and ventral (B); ETVP 3293 in dorsal (C) and ventral (D); ETVP 3306 in dorsal (E) and ventral (F). Anterior is to the top of the page. Scale bar = 1 mm.

## Descriptions

The description of these disarticulated cranial elements can act as a foundation for a more in-depth study on ontogenetic variation. Using the descriptions, geometric morphometrics can be used to map the changes through ontogeny and potentially demonstrate a greater degree of change. These descriptions can also allow for comparisons with other species of the genus to document how certain structures differ between species, and possibly identify new diagnostic characteristics.

The cranial elements of Crotalinae species have played a role in many studies, and with an in-depth description of disarticulated elements, *T. wagleri* could be used in more studies to better understand the relationships within Asian pitvipers. Diagnostic characteristics of certain taxa have been previously identified [9, 18]; thus with the addition of full descriptions of disarticulated elements, additional characteristics could be identified. While ontogeny was minimally approached here, the descriptions produced can allow comparisons of different age groups so that ontogeny could be better understood; since *T. wagleri* exhibits such drastic color changes in ontogeny [3]. *Tropidolaemus wagleri* could also add to the understanding of pitviper evolution. Many authors have commented on the basal nature of *T. wagleri* within Crotalinae. With an in-depth description of skull morphology the evolution and development of certain characters could be mapped to estimate the sequence of evolution for pitvipers [3, 9, 12, 22, 23, 25].

Many previous studies observing the cranial osteology of members of the Crotalinae compared the cranial characteristics to other specimens to hypothesize phylogeny, evolution, taxonomy, and zoogeography. Among the previous studies observing cranial elements some described only the differences/similarities among studied taxa, while others offered more detailed descriptions. The study here provides a detailed description of multiple elements without comparison to other taxa. Brattstrom [9] did reference ontogeny within the Crotalinae, stating that he did not observe major shape change after birth with the exception of *Lachesis muta*.

Guo and Zhao [18] identified characteristics of individual skull elements that can be used to identify and differentiate between five genera and nine species of Asian pitvipers. Using the characters in Guo and Zhao [18] *Tropidolaemus wagleri* appears to possess the following character states: Border of maxillary cavity: projection; Parietal ridge: strong; Parietal shape; T-shaped; Palatine shape: crescent and forked; Ectopterygoid anterior lateral process: broad.

Guo et al. [21] identified 12 osteological characteristics in order to hypothesize the evolution of Asian pitvipers. No specimens of *Tropidolaemus spp* were used in the study. Using the 12 characters described in Guo et al. [21] *Tropidolaemus wagleri* appears to possess the following character states: Dorsal process of dentary does not reach the ventral process; the shape of the anterior end of the ectopterygoid is broad; thin flap on the posterior ventrolateral edge of the lower jaw is present; the shape of the palatine is forked; projection on border of maxillary cavity present; shape of the parietal ridge is T-shaped; pterygoid teeth extend to posterior margin of articulation with ectopterygoid; squamosal does extend beyond the posterior end of the braincase; splenial and angular are separated; maxillary cavity is closed.

Previous studies have focused on taxonomy, soft anatomy, molecular sequencing, and brief osteologic comparisons and identifying characters of *Tropidolaemus wagleri*. The study here is the logical next step from previous studies and opens the door for studies on identifying isolated elements and understanding full ontogenetic change within one taxon, taking the next step toward paleontological studies.

## Conclusion

- *Tropidolaemus wagleri* has been focused on, mostly in a taxonomic sense and did not provide data to be utilized for paleontological finds, hence this study.
- Description of cranial elements allows for identification of disarticulated elements in fossilized specimens.
- Brattstrom noticed little ontogenetic change in crotalids with the exception of *Lachesis muta*. Brattstrom observed “There is no major change in the shape of the various bones of the skulls of crotalids after birth” [9:190]. Contra Brattstrom [9], my study shows the greatest change occurs early on with the most dramatic change between Age Ranks 1 and 2.
- The bones that exhibit the most ontogenetic change in this study are the maxilla, parietal, and prefrontal.
- Most ontogenetic changes appear to be the result of the isometric growth and the “filling out” of the bones’ shape.

## Abbreviations

AR: Age Rank
APPSU: Appalachian State University
ASU: Appalachian State University
ETVP: East Tennessee State University Vertebrate Paleontology Collections
SVL: snout-vent length.

